# The cell cycle regulator Cyclin E1 drives macrophage-associated inflammatory responses in acute immune-mediated hepatitis

**DOI:** 10.64898/2026.09.30.755646

**Authors:** Christian Penners, Cheng Lin, Alexander M. Jans, Antje Mohs, Frank Tacke, Matthias Bartneck, Roland Sonntag, Christian Liedtke

## Abstract

**Background and aims:** Cyclin E1 is a canonical regulator of cell cycle progression known to promote liver fibrosis and hepatocarcinogenesis. However, its precise contribution to hepatic immune responses remains poorly understood. We investigated whether Cyclin E1 regulates macrophage-associated inflammation and acute immune-mediated liver injury.

**Methods:** Hepatic immune populations were analyzed in untreated and concanavalin A-treated wildtype and constitutive *Ccne1*^−/−^ mice by flow cytometry. Liver injury was assessed by survival, histopathology, and caspase activation. Inflammatory mediators were quantified by qPCR and ELISA. Bone marrow-derived macrophages were examined during differentiation and pro-inflammatory polarization using flow cytometry, cytokine measurements, and RNA-sequencing. Lipid nanoparticle-mediated delivery of *Ccne1*-specific siRNA was evaluated in J774A.1 macrophage-like cells.

**Results:** Under homeostatic conditions, *Ccne1*^−/−^ mice exhibited reduced hepatic immune cell numbers, including Kupffer cells and CD4^+^ and CD8^+^ lymphoid cells. Following concanavalin A administration, *Ccne1*^−/−^ mice showed improved survival, reduced hepatic necrosis, and diminished caspase-3 and caspase-8 activation. Protection was associated with reduced Kupffer and CD8^+^ lymphoid cell abundance and attenuated inflammatory mediator expression. Cyclin E1 deficiency did not substantially alter bone marrow-derived macrophage differentiation or cell cycle distribution. However, pro-inflammatory *Ccne1*^−/−^ macrophages showed reduced interleukin-6 expression, attenuated inflammatory gene programs, and lower inferred IRF1 and NF-κB1 activity. *Ccne1* knockdown similarly reduced interleukin-6 expression in pro-inflammatory J774A.1 cells.

**Conclusions:** Cyclin E1 contributes to hepatic immune homeostasis and promotes macrophage-associated inflammatory signaling during immune-mediated hepatitis. Its inflammatory function is uncoupled from overt changes in macrophage cell cycle progression and may involve regulation of IRF1- and NF-κB-associated transcriptional programs. Targeting Cyclin E1 may therefore provide a strategy to attenuate selected inflammatory responses in liver disease.

## Introduction

Acute and chronic liver injury are shaped by complex interactions between parenchymal liver cells and non-parenchymal immune cell populations. In particular, hepatic macrophages are central regulators of liver inflammation, tissue injury, and resolution. The liver contains resident macrophages, Kupffer cells, as well as recruited monocyte-derived macrophages, which accumulate during inflammatory liver damage^1^. Depending on the local microenvironment, these macrophage populations can promote inflammation through secretion of cytokines and chemokines such as interleukin 6 (IL6), tumor necrosis factor (TNF), chemokine (C-C motif) ligand 2 (CCL2), and CCL5, but may also contribute to immune regulation through anti-inflammatory mediators including interleukin 10 (IL10)^2, 3^. Thus, macrophage plasticity is a critical determinant of hepatic immune homeostasis and inflammatory liver disease.

Concanavalin A (ConA)-induced hepatitis is a well-established model of acute immune-mediated liver injury that recapitulates key features of autoimmune-like liver inflammation^4^ including the release of inflammatory mediators such as TNF and interferon gamma (IFNγ) by activated T cells and subsequent hepatocellular injury and necrosis^5–9^. Macrophage-derived cytokines and chemokines amplify the inflammatory response and contribute to monocyte recruitment and macrophage accumulation at sites of tissue injury. Therefore, the ConA model provides a suitable experimental system to dissect immune cell-driven pathways in acute liver damage.

E-type cyclins are classically recognized as regulators of G1/S-phase transition through activation of cyclin-dependent kinase 2 (CDK2) and downstream E2F-dependent transcriptional programs^10^. However, genetic studies demonstrated that mice lacking either Cyclin E1 encoded by the *Ccne1* or Cyclin E2 encoded by the *Ccne2* gene are viable, indicating that individual E-type cyclins are not universally required for cell cycle progression in all tissues^11, 12^. In the liver, Cyclin E1 has emerged as a disease-relevant regulator beyond its canonical role in cell cycle control. Previous studies showed that Cyclin E1 deficiency does not affect liver regeneration, but protects from experimental liver fibrosis, and impairs hepatocarcinogenesis, whereas Cyclin E2 can exert distinct and partly opposing functions^13–16^.

Moreover, therapeutic systemic inhibition of *Ccne1* by siRNA delivery attenuated liver fibrosis and inflammation in experimental liver injury models^17^, suggesting that Cyclin E1 may represent a druggable regulator of liver pathology.

Although these studies point to a disease-promoting role of Cyclin E1 in the liver, the underlying cell type-specific mechanisms remain incompletely understood. Notably, Cyclin E1 inhibition in liver tumors was associated with reduced expression of inflammatory mediators, including IL6^16^, raising the possibility that Cyclin E1 may regulate liver disease not only through effects on proliferating parenchymal or mesenchymal cells, but also by modulating immune cell function. This concept is further supported by evidence for tissue-specific Cyclin E1 interaction networks, suggesting context-dependent and potentially non-canonical functions of Cyclin E1^18^.

Here, we investigated whether Cyclin E1 controls hepatic immune responses with a particular focus on macrophages. Using constitutive *Ccne1* knockout mice, the ConA model of acute immune-mediated hepatitis, primary bone marrow-derived macrophages, and lipid nanoparticle-mediated siRNA delivery, we analyzed the role of Cyclin E1 in hepatic immune cell composition, macrophage differentiation and polarization, and inflammatory cytokine production *in vivo* and *in vitro*. We will provide evidence that Cyclin E1 can act not only as a cell cycle regulator, but also as a cell cycle-independent pro-inflammatory factor in macrophages.

## Results

We determined whether Cyclin E1 contributes to hepatic immune cell homeostasis. To this end, we enzymatically digested livers from untreated wildtype and *Ccne1*^−/−^ mice and analyzed isolated immune cells by flow cytometry. Using a myeloid gating strategy (Supplementary Figure 1A), *Ccne1*^−/−^ mice exhibited a reduced number of total CD45^+^ hepatic immune cells compared to wildtype controls. Analysis of specific subpopulations revealed no significant differences in monocyte or monocyte-derived macrophage (MoMF) numbers, whereas Kupffer cell abundance was significantly decreased in the absence of Cyclin E1 (Figure 1A). In addition to myeloid populations, lymphoid cells were analyzed using a different gating strategy (Supplementary Figure 1B, Figure 1B). Consistent with the findings above, *Ccne1*^−/−^ mice showed an overall reduction in total hepatic immune cell numbers. More specifically, this decrease was associated with reduced abundance of both CD4^+^ and CD8^+^ T cells. Collectively, these data indicate that Cyclin E1 plays a critical role in hepatic immune cell homeostasis.

**Figure 1:**
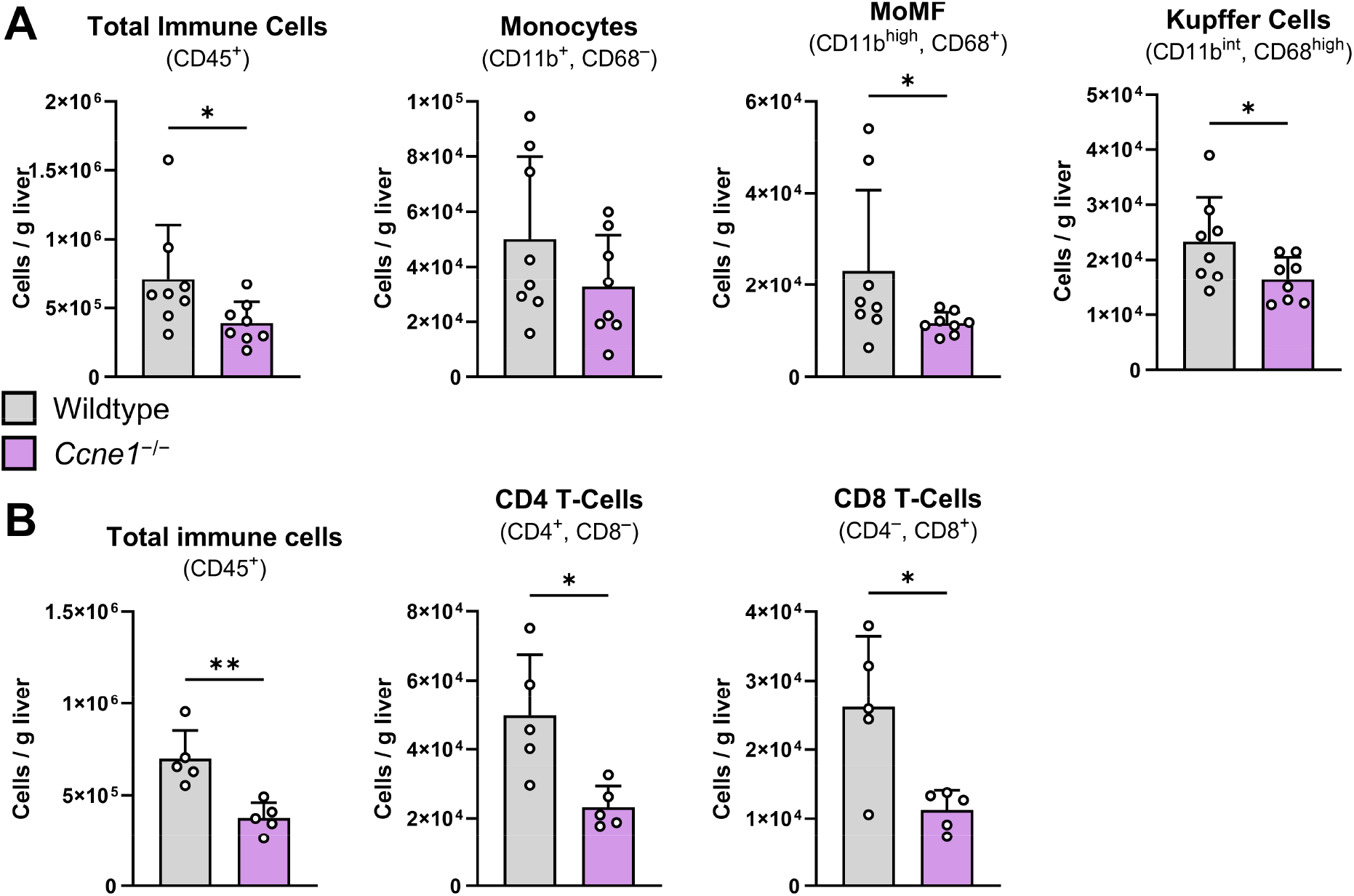
Cyclin E1 modulates the hepatic immune cell landscape in mice. Livers of 8-10-week-old wildtype (n = 8) and *Ccne1*^−/−^ (n = 8) mice were analyzed for their hepatic immune cell composition by flow cytometry. APC-labeled beads were added to a defined mass of liver tissue as an internal control to calculate immune cells per gram liver tissue. **(A)** Analysis of hepatic immune cells from the myeloid lineage and **(B)** from the lymphoid lineage. Data is shown as mean ± SD. *: *p* ≤ 0.05, **: *p* ≤ 0.01.

To investigate the functional consequences of Cyclin E1 deficiency under inflammatory conditions and to identify potential immune effector populations, we employed the Concanavalin A (ConA)-induced hepatitis model. This well-established model recapitulates key features of acute immune-mediated liver injury in humans and is characterized by robust immune activation^4, 7^. ConA-induced liver damage is primarily driven by activated T cells, with additional contributions from macrophages that amplify inflammatory responses.

Wildtype and *Ccne1*^−/−^ mice were challenged with ConA and analyzed after 24 h (Figure 2A). Liver-to-body weight ratios were comparable between genotypes, indicating no gross differences in liver mass following treatment (Figure 2B). Notably, while only 37.5 % of wildtype mice survived ConA administration at the applied dose, all *Ccne1*^−/−^ mice were protected from treatment-induced mortality (Figure 2C). To assess the extent of liver injury in surviving animals, histopathological analyses were performed on H&E-stained liver sections. As expected, ConA treatment induced tissue necrosis in both groups. However, quantitative evaluation revealed significantly greater necrotic areas in wildtype mice (53 ± 17 %) compared to *Ccne1*^−/−^ mice (16 ± 9 %) (Figure 2D), indicating markedly attenuated liver damage in the absence of Cyclin E1. Western blot analysis of liver lysates further revealed increased levels of cleaved caspase-8 and cleaved caspase-3 in ConA-treated wildtype mice compared to *Ccne1*^−/−^ mice and untreated controls, indicating enhanced activation of apoptosis-associated cell death pathways in wildtype livers (Figure 2E). Collectively, these data demonstrate that Cyclin E1 deficiency protects mice from ConA-induced acute liver injury, as reflected by improved survival, reduced tissue necrosis, and diminished activation of apoptosis-associated cell death pathways.

**Figure 2:**
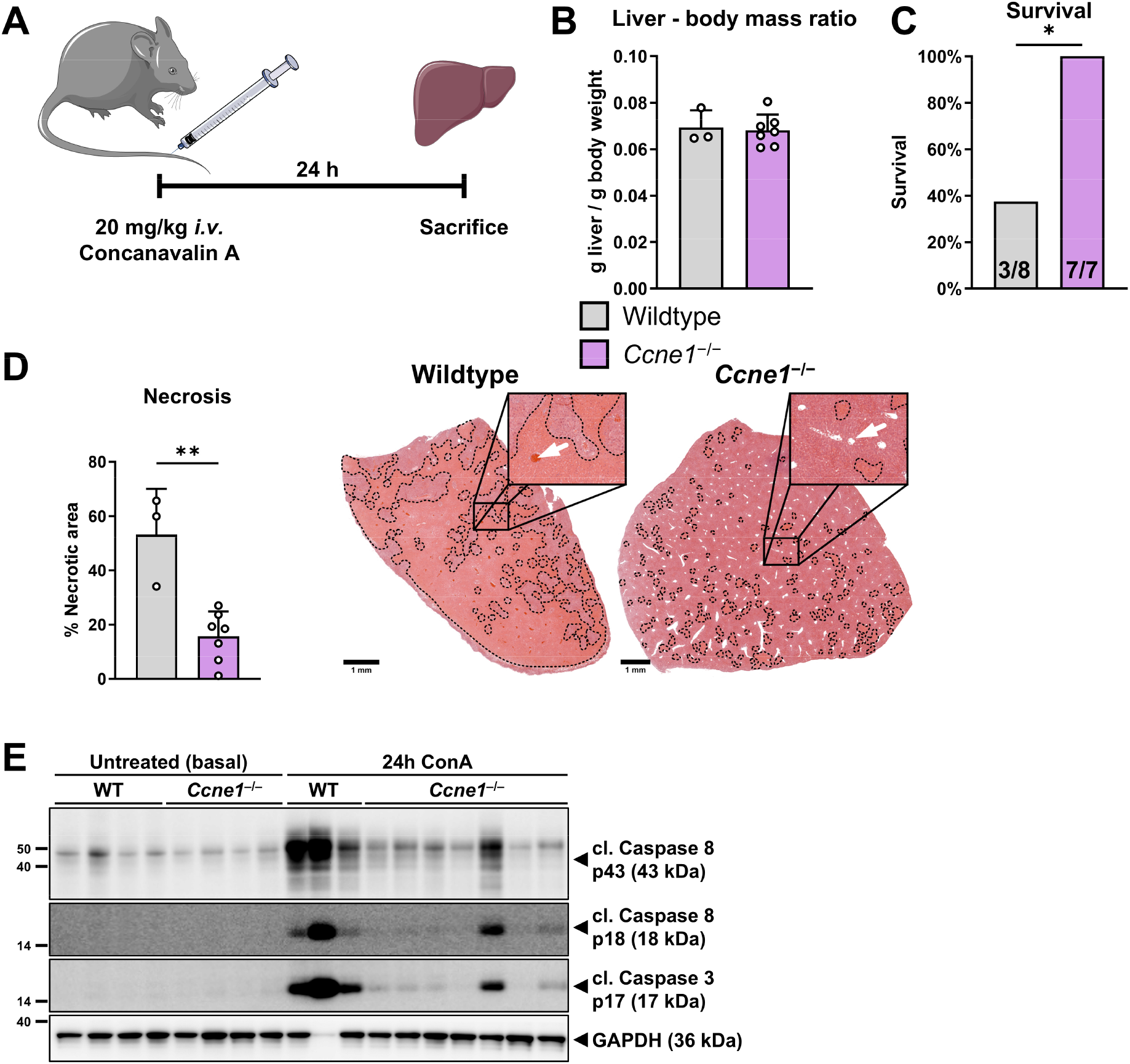
Cyclin E1-deficient mice are protected from ConA-mediated liver injury and mortality. **(A)** Wildtype (n = 8) and *Ccne1*^−/−^ (n = 7) mice received an intravenous injection of 20 mg/kg ConA and their livers were analyzed after 24 h. **(B)** Liver mass index after ConA treatment of surviving wildtype (n = 3) and *Ccne1*^−/−^ (n = 7) mice calculated as liver to body mass ratio. **(C)** Survival analysis of ConA-treated mice shown as percent survival with absolute numbers shown within respective bars. **(D)** Quantification of necrotic liver tissue area in percent as quantified from H&E-stained livers as shown in representative image on the right. Necrotic areas are highlighted by dashed lines and appear slightly orange. White arrows in enlarged views indicate blood vessels, which are characterized by intravascular hemorrhage in wildtype livers. **(E)** Western blot from liver lysates of untreated wildtype (n = 4) and *Ccne1*^−/−^ (n = 4) mice and ConA-treated wildtype (n = 3) and *Ccne1*^−/−^ (n = 7) mice showing protein levels of cleaved caspase-3 (p17 fragment) and cleaved caspase-8 (p43 and p18 fragment). GAPDH expression is shown as reference. Data is shown as mean ± SD. *: *p* ≤ 0.05, **: *p* ≤ 0.01.

Given that ConA-induced pathology is largely immune-mediated, hepatic immune cell composition in surviving mice was analyzed by flow cytometry. While total immune cells, monocyte and monocyte-derived macrophage (MoMF) populations were comparable between genotypes, Kupffer cell numbers were significantly reduced (> 50 %) in *Ccne1*^−/−^ mice compared to wildtype controls (Figure 3A). Analysis of lymphoid populations revealed significantly reduced CD8^+^ T-cell numbers in *Ccne1*^−/−^ livers, while CD4^+^ T cells and regulatory T cells were not significantly altered (Figure 3B). To determine whether these changes were associated with altered inflammatory signaling, cytokine and chemokine expression were quantified in liver tissue from wildtype and *Ccne1*^−/−^ mice under basal conditions and following ConA treatment by qPCR and ELISA (Figure 3C+D). In wildtype livers, ConA administration induced robust expression of interleukin-6 (*Il6*/IL6), interferon gamma (*Ifng*/IFNG), and interleukin-10 (*Il10*/IL10) at both the mRNA and protein level. This induction was significantly attenuated in *Ccne1*^−/−^ mice. Similarly, tumor necrosis factor alpha (*Tnf*/TNF) and chemokine (C-C motif) ligand 2 (*Ccl2*/CCL2) were increased at the protein level in wildtype livers following ConA challenge, whereas these responses were significantly reduced in Cyclin E1-deficient tissues, despite unchanged mRNA expression. Together, these findings indicate that Cyclin E1 deficiency attenuates ConA-induced immune activation and is associated with reduced Kupffer cell abundance and diminished expression of macrophage-associated cytokines and chemokines.

**Figure 3:**
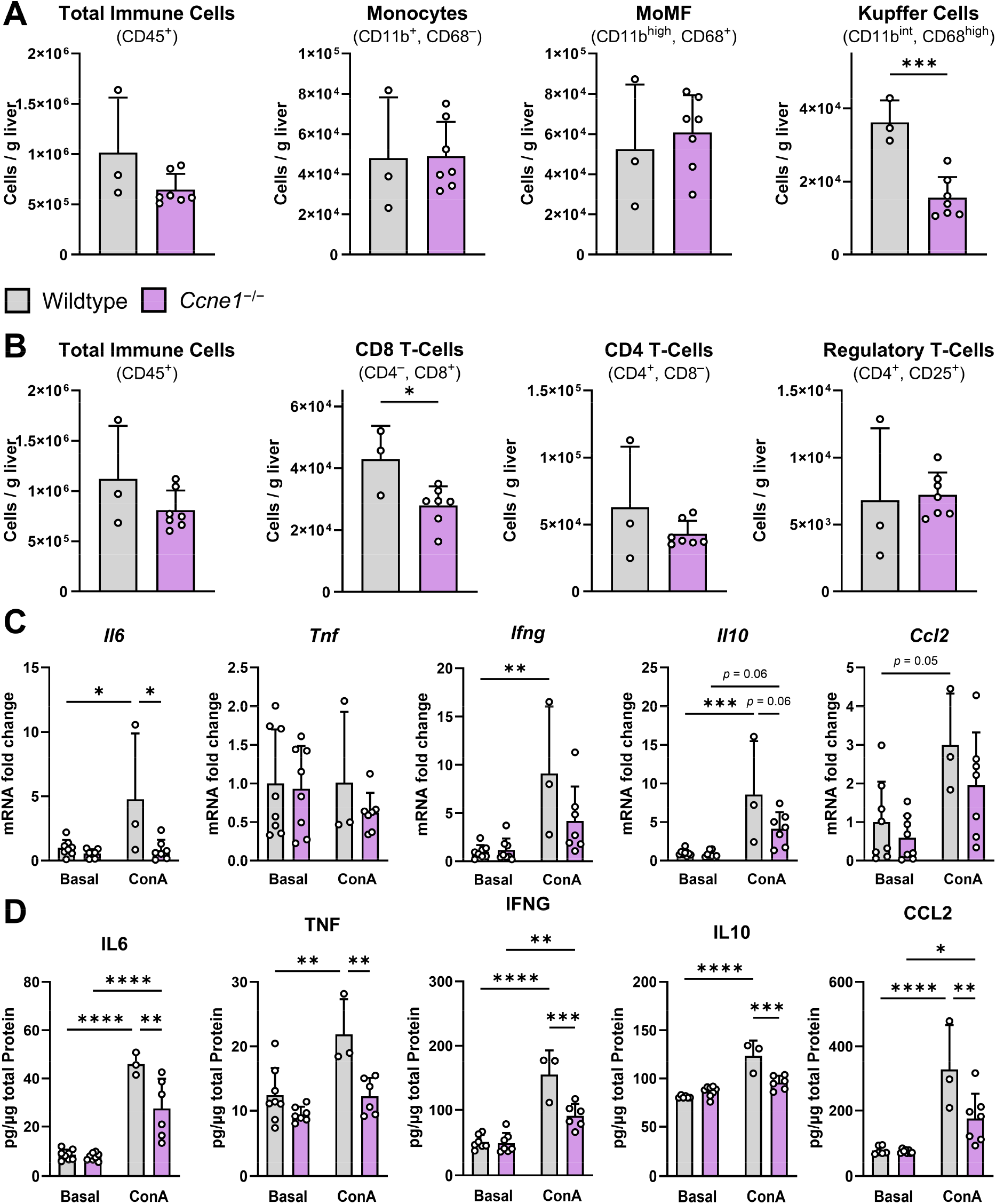
Cyclin E1 modulates immune cell composition and inflammatory response in ConA-mediated hepatitis. Wildtype (n = 3) and *Ccne1*^−/−^ (n = 7) mice were treated with ConA as described in Figure 2A followed by liver dissection and analysis of (**A**) myeloid and (**B**) lymphoid immune cell composition by flow cytometry. APC-labeled beads were added to a defined mass of liver tissue as an internal control to calculate immune cells per gram liver tissue. Total mRNA and protein extracts were isolated from ConA-treated mice and untreated wildtypes (n = 8) and *Ccne1*^−/−^ mice (n = 8). (**C**) Relative gene expression levels of inflammation-associated genes were analyzed by qPCR. Gene expression was normalized to *Actb* expression and calculated as fold change compared to untreated wildtype mice. (**D**) Corresponding tissue protein levels were determined by ELISA using liver protein isolates. Concentrations were normalized to the concentration of total protein in liver lysates as determined by Bradford assay. Data is shown as mean ± SD. *: *p* ≤ 0.05, **: *p* ≤ 0.01, ***: *p* ≤ 0.001.

The reduced Kupffer cell abundance and attenuated macrophage-associated inflammatory mediator expression observed in *Ccne1*^−/−^ livers suggested that Cyclin E1 may regulate macrophage biology. We therefore investigated whether Cyclin E1 affects macrophage differentiation and cell cycle progression *in vitro*. Bone marrow precursor cells from wildtype and *Ccne1*^−/−^ mice were isolated and differentiated into bone marrow-derived macrophages (BMDMs). Differentiation kinetics were monitored over a period of 7 days by flow cytometry using the markers CD45, CD11b, and CD68 to distinguish myeloid subpopulations (Figure 4A). Quantitative analysis of precursor cells, monocytes, and macrophages revealed no significant differences between wildtype and *Ccne1*^−/−^ cells throughout the observation period (Figure 4B), indicating that Cyclin E1 is dispensable for macrophage differentiation. To assess potential effects on cell cycle activity during differentiation, DNA content was analyzed by DAPI staining and subsequent flow cytometry (Figure 4C). BMDMs exhibited a transient increase in S-phase activity on day 3, followed by a peak in G2/M-phase on day 4. Across most time points, cell cycle distribution was comparable between genotypes, with no differences observed in G0/G1 or S phase. A modest reduction in the proportion of *Ccne1*^−/−^ cells in G2/M phase was detected on day 6. Overall, these data indicate that loss of Cyclin E1 does not substantially affect cell cycle progression or differentiation of bone marrow-derived macrophages *in vitro*. Previous studies have suggested that loss of Cyclin E1 can be compensated by other cyclins, including Cyclin E2 as well as A- and D-type cyclins^19^. To determine whether compensatory mechanisms are activated upon *Ccne1* deletion, the expression of key cell cycle regulators was analyzed in bone marrow-derived cells by qPCR (Figure 4D). As expected, *Ccne1* transcripts were undetectable in *Ccne1*^−/−^ cells. Importantly, the expression profiles of other cell cycle-associated genes (*i.e. Ccne2*, *Cdk2*, *Ccna2*, *Ccnb1*, *Ccnd1*) remained largely unchanged in the absence of Cyclin E1, arguing against compensatory upregulation.

**Figure 4:**
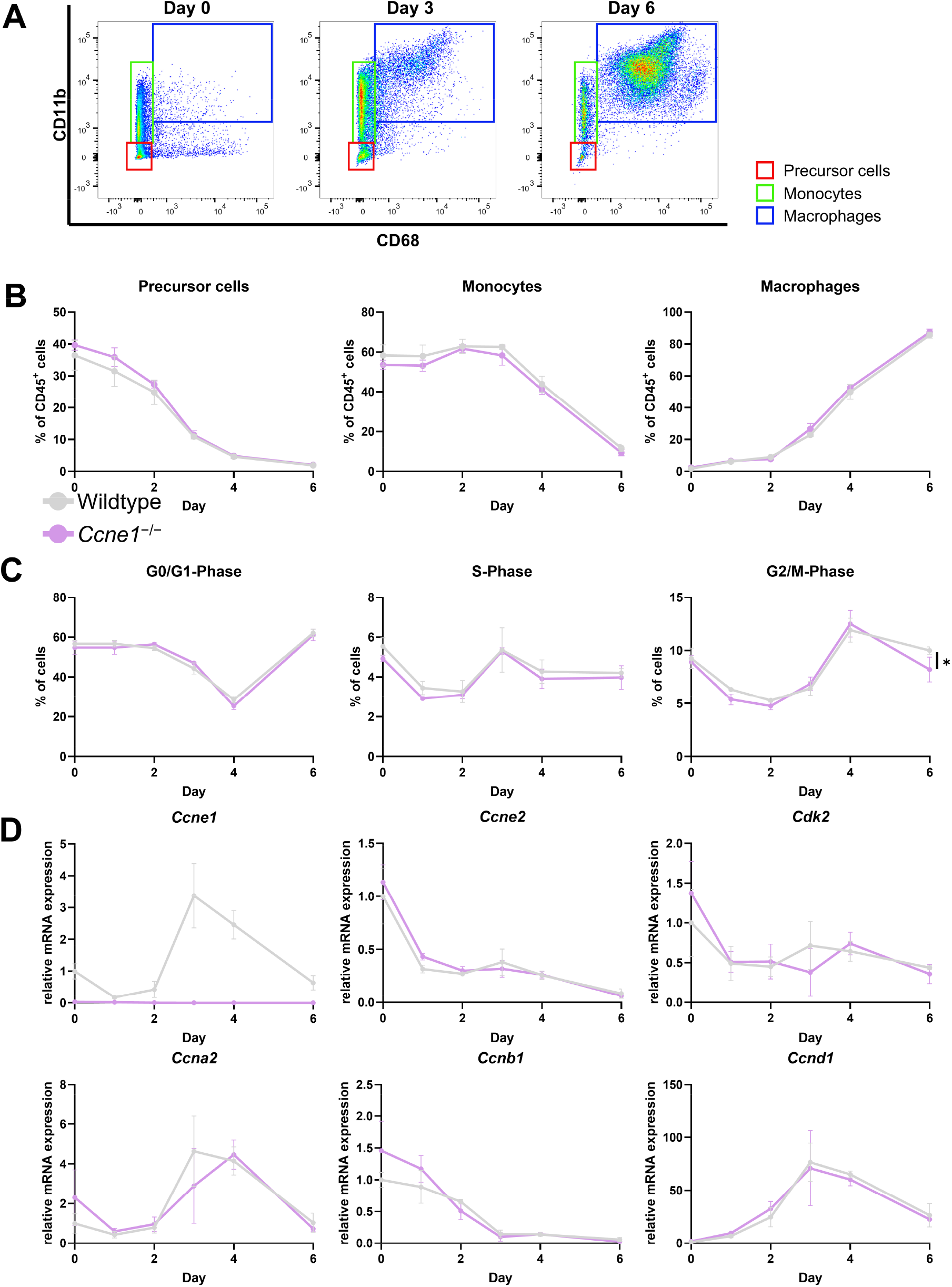
Loss of Cyclin E1 does not affect macrophage differentiation from precursors. Wildtype (n = 3) and *Ccne1*^−/−^ (n = 3) mice were sacrificed to isolate bone marrow for *in vitro* differentiation of bone marrow derived macrophages. Cells were cultured in presence of L929 cell supernatants which leads to macrophage differentiation from precursor cells. (**A**) Differentiation was monitored by flow cytometry on six consecutive days (except day 5) using antibodies against CD45, CD11b and CD68 to distinguish between precursors, monocytes and macrophages as indicated by highlighted gating. (**B**) Subpopulations were quantified over time relative to the respective number of CD45^+^ cells. (**C**) Cell cycle phase distribution during differentiation was determined for all cells by flow cytometry and DNA content measurement using DAPI. Cell numbers in each cell cycle phase are shown as percentage relative to the total cell number. (**D**) Total RNA was isolated at each timepoint, and relative mRNA expression of cell cycle mediators was analyzed by qPCR. Gene expression was normalized to *Actb* expression and calculated as fold change compared to wildtype cells at day 0. Data is shown as mean ± SD. *: *p* ≤ 0.05.

Taking together, these data indicate that Cyclin E1 deficiency neither induces compensatory expression of other cyclins or CDKs nor alters the temporal dynamics of cell cycle progression during macrophage differentiation.

To determine whether Cyclin E1 influences inflammatory macrophage polarization or associated cell cycle activity, fully differentiated BMDMs from wildtype and *Ccne1*^−/−^ mice were polarized *in vitro* towards classically activated macrophages referred to as M1 macrophages or left unstimulated (M0) (Figure 5A). Successful stimulation was confirmed by qPCR analysis of *Nos2* expression, a well-established marker for M1 polarization (Figure 5B). As expected, *Nos2* expression was strongly induced in M1-polarized macrophages and comparable between wildtype and *Ccne1*^−/−^ cells, indicating that Cyclin E1 is not required for macrophage polarization. To assess potential effects on cell cycle activity, DNA content was analyzed by flow cytometry in M0 and M1 macrophages (Figure 5C). No significant differences between genotypes were observed across all polarization states, and the proportion of cells in G0/G1 phase remained unchanged. In wildtype cells, M1 polarization was associated with a modest decrease in S-phase cells compared to M0. However, these effects were minor, suggesting that polarization exerts only limited influence on cell cycle distribution. Overall, these data indicate that Cyclin E1 is dispensable for macrophage polarization and has no major impact on cell cycle phase distribution under these conditions. Next, the expression of key cell cycle regulators was assessed in polarized BMDMs by qPCR (Figure 5D). No significant differences were observed between wildtype and *Ccne1*^−/−^ cells across both polarization conditions. Polarization itself, however, influenced the expression of specific cell cycle genes. In M1-polarized BMDMs, transcript levels of *Ccne2* and *Ccna2* were significantly reduced compared to unstimulated (M0) cells. A similar reduction was observed for *Ccnb1* and *Ccnd1*, which was only significant in wildtype cells but with a clear comparable trend in *Ccne1*^−/−^ cells. Of note, *Ccne1* expression was not downregulated in M1-polarized wildtype macrophages. Macrophage M1-polarization is associated with the production and secretion of cytokines. To assess the impact of Cyclin E1 deficiency on this process, mRNA expression and protein levels of key inflammatory cytokines (*Tnf*/TNF, *Il6*/IL6, and *Il10*/IL10) were measured in M1-polarized BMDMs by qPCR and ELISA (Figure 5E+F). In wildtype BMDMs, M1 polarization led to robust induction of *Tnf*/TNF and *Il6*/IL6 at both the transcript and protein levels compared to unstimulated (M0) cells. *Il10*/IL10 levels were also increased under M1 conditions, however, this reached only statistical significance in *Ccne1*^−/−^ cells. Notably, Cyclin E1 deficiency significantly altered this cytokine profile: *Il10* mRNA expression showed a trend towards increased expression (*p* = 0.08), whereas *Il6*/IL6 expression was significantly reduced at both mRNA and protein levels in *Ccne1*^−/−^ M1-polarized macrophages. These findings indicate that Cyclin E1 modulates cytokine production in M1 macrophages, particularly influencing the balance between pro- and anti-inflammatory mediators. Corresponding analyses were also performed in M2a-polarized BMDMs and revealed only minor genotype-dependent differences, none of which reached statistical significance (Supplementary Figure 2).

**Figure 5:**
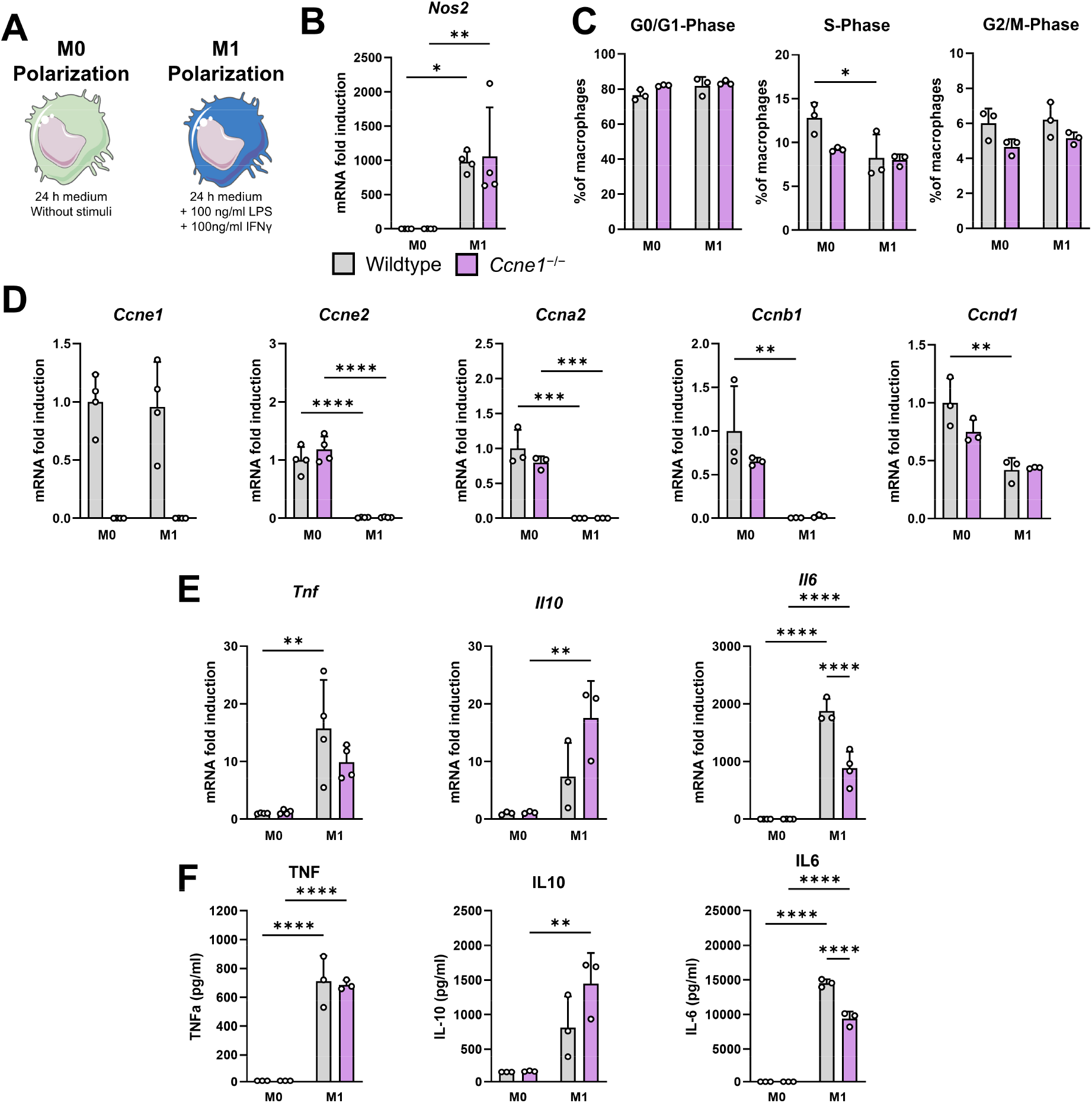
Cyclin E1-deficient macrophages show reduced pro-inflammatory properties upon M1 polarization. Wildtype (n = 3 - 4) and *Ccne1*^−/−^ (n = 3 - 4) mice were sacrificed to isolate bone marrow for *in vitro* generation of BMDMs followed by their pro-inflammatory polarization (M1) or were left untreated (M0). (**A**) Schematic representation of M0 or M1 polarization conditions. (**B**) Relative *Nos2* gene expression was determined by qPCR. Expression was normalized to *Actb* levels and calculated as fold change compared to M0-polarized wildtype BMDMs. (**C**) Cell cycle phases were determined by flow cytometry using DAPI to measure cellular DNA content. The cell numbers in each phase are shown relative to all analyzed BMDMs. Gene expression of cell cycle-related genes (**D**) and inflammatory mediators (**E**) was determined by qPCR. Expression was normalized to *Actb* levels and calculated as fold change compared to M0-polarized wildtype BMDMs. (**F**) Protein levels of inflammatory mediators in BMDM supernatants were measured by ELISA. Data is shown as mean ± SD. *: *p* ≤ 0.05, **: *p* ≤ 0.01, ***: *p* ≤ 0.001, ****: *p* ≤ 0.0001.

To obtain a broader view of transcriptional programs associated with macrophage polarization and Cyclin E1 deficiency, bulk RNA sequencing (RNA-seq) was performed on M0 and M1-polarized wildtype and *Ccne1*^−/−^ BMDMs. As expected, M1 polarization induced a pronounced transcriptional shift in wildtype cells, with approximately 50% of detected transcripts being differentially expressed (Supplementary Figure 3A+B). Overrepresentation analysis (ORA) of genes upregulated in M1-polarized wildtype BMDMs revealed significant enrichment of gene ontology terms related to inflammation and macrophage activation (Figure 6A). In contrast, downregulated genes were strongly enriched for cell cycle–associated processes, particularly those linked to S-phase progression and mitosis. Consistently, multiple chemokines and cytokines as well as NF-κB related genes were induced, whereas genes related to the origin recognition complex, cyclins and cyclin-dependent kinases were suppressed (Figure 6B). Notably, *Ccne1* expression did not follow this general downregulation pattern, consistent with the qPCR data (compare Figure 5D). Transcription factor activity inference analysis further indicated increased activity of inflammation-associated regulators, including IRF1 and NF-κB1, while activities of E2F family transcription factors, which are central to cell cycle control, were reduced in M1-polarized BMDMs (Figure 6C). Overall comparable results were observed when comparing M1-polarized *Ccne1*^−/−^ BMDMs to M0-polarized ones (Supplementary Figure 3C-G).

**Figure 6:**
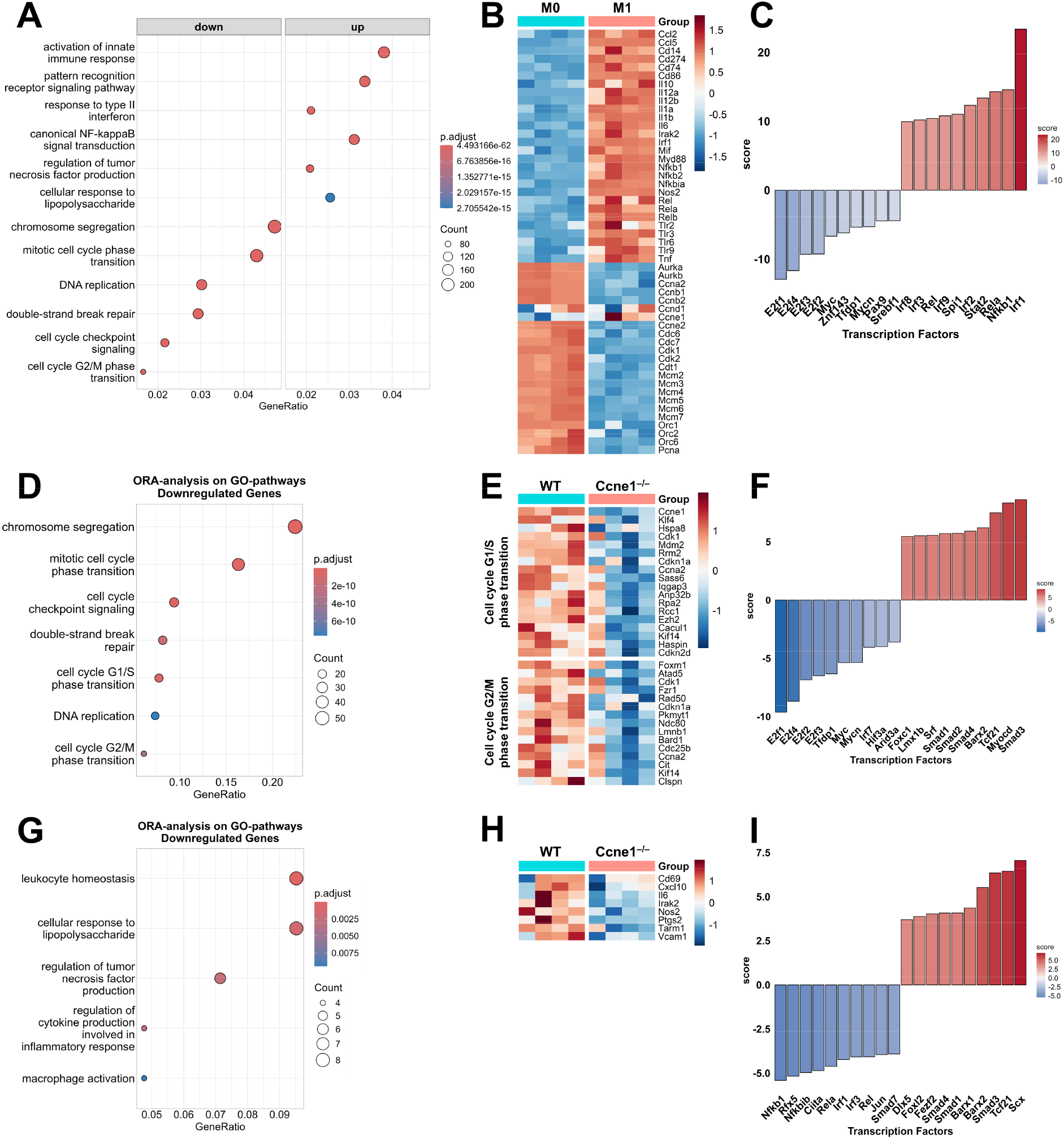
Cyclin E1 modulates basal and pro-inflammatory gene regulation in BMDMs. Wildtype (n = 4) and *Ccne1*^−/−^ (n = 4) mice were sacrificed to isolate bone marrow for *in vitro* generation of BMDMs followed by their pro-inflammatory polarization (M1) or were left untreated (M0) as described in Figure 5A. Total mRNA was isolated and bulk RNA-seq was performed. (**A**) Differential gene expression between M1-polarized wildtype BMDMs compared to M0 wildtype BMDMs was analyzed. Over representation analysis was performed on up- or down-regulated genes using gene ontology pathways. Results of selected significant deregulated pathways are shown as dot plot indicating number of regulated genes by size and statistical significance by color. (**B**) Heatmap of selected genes involved in regulation of inflammation or cell cycle. (**C**) Transcription factor activity scores were calculated using transcription factor activity inference. The ten statistically significant most active or deactivated transcription factors are shown as bar plot. (**D**) Differential gene expression between M0 BMDMs from *Ccne1*^−/−^ mice compared to wildtype ones was analyzed. Over representation analysis was performed on down-regulated genes using gene ontology pathways. Results of selected significant deregulated pathways are shown as dot plot. (**E**) Heatmap showing deregulated genes belonging to the indicated gene ontology pathways. (**F**) Results of transcription factor activity inference analysis. (**G**) Differential gene expression between M1 BMDMs from *Ccne1*^−/−^ mice compared to wildtype ones was analyzed. Over representation analysis was performed on down-regulated genes using gene ontology pathways. Results of selected significant deregulated pathways are shown as dot plot. (**H**) Heatmap showing deregulated pro-inflammatory genes. (**I**) Results of transcription factor activity inference analysis.

Next, the impact of Cyclin E1 deletion on M0-polarized BMDMs was examined in greater detail. M0 *Ccne1*^−/−^ BMDMs showed a slightly different expression profile and in total 434 differentially expressed genes compared to wildtype controls (Supplementary Figure 4A+B). Contrary to the qPCR and flow cytometry data, which suggested no impact on cell cycle activity, ORA analysis on differentially expressed genes indicates deregulated gene expression of cell cycle-related genes (Figure 6D). However, the cell cycle-related genes identified by RNA-seq only partially overlapped with the genes analyzed by qPCR. Among the qPCR-tested regulators, only *Ccna2* was included in the differentially expressed genes associated with G1/S or G2/M phase transition (Figure 6E). Nevertheless, transcription factor activity inference supported a reduced cell cycle-associated transcriptional program in unstimulated *Ccne1*^−/−^ BMDMs, as the four most strongly deactivated transcription factors belonged to the E2F family, which is closely linked to cell cycle regulation (Figure 6F).

We next compared M1-polarized *Ccne1*^−/−^ and wildtype BMDMs directly. Although global transcriptional differences between both genotypes were relatively modest (Supplementary Figure 4B+C), the RNA-seq data supported the reduced inflammatory phenotype observed by qPCR and ELISA. ORA analysis of genes downregulated in *Ccne1*^−/−^ BMDMs revealed significant enrichment of pathways related to inflammatory macrophage activation and polarization (Figure 6G). Besides others, *Il6* was among the mediators for which reduced expression has been observed (Figure 6H). In line with these findings, transcription factor activity inference analysis predicted reduced activity of pro-inflammatory transcription factors, including IRF1 and NF-κB1, in M1-polarized *Ccne1*^−/−^ BMDMs (Figure 6I).

In summary, RNA-seq data show reduced cell cycle-related gene expression in M0 *Ccne1*^−/−^ BMDMs compared to wildtype controls. M1 polarization of BMDMs is, as expected, associated with the expression of pro-inflammatory mediators and further strong downregulation of several cell cycle-related genes but not Cyclin E1. Lack of Cyclin E1 in M1-polarized BMDMs, however, leads to reduced expression of pro-inflammatory genes including lower activity of inflammation driving transcription factors.

The preceding data suggested that Cyclin E1 inhibition may promote anti-inflammatory features in M1-polarized macrophages, reflected by reduced expression of pro-inflammatory mediators including *Il6*. We therefore next investigated whether this observation could be translated into a potential therapeutic approach using RNA interference. Our group has previously shown that siRNA-mediated inhibition of Cyclin E1 in the murine liver is technically feasible and sufficient to attenuate liver fibrosis progression^17^. Based on these findings, we assessed whether siRNA-mediated knockdown of Cyclin E1 could suppress pro-inflammatory properties in the murine macrophage cell line J774A.1. First, we evaluated whether J774A.1 cells are capable of taking up siRNA-loaded lipid nanoparticles (LNPs). Cells were pre-incubated with LNPs loaded with either a non-targeting scrambled control siRNA, hereafter referred to as control siRNA, or TEX615 fluorophore-labeled siRNA, followed by M0 or M1 polarization (Figure 7A). LNP uptake was assessed by fluorescence microscopy and flow cytometry (Figure 7B+C). Both approaches demonstrated efficient uptake of siRNA-loaded LNPs by J774A.1 macrophages. Flow cytometric analysis further revealed that nearly all J774A.1 cells incorporated LNPs under both M0 and M1-polarized conditions making them a suitable *in vitro* model to evaluate siRNA-mediated Cyclin E1 inhibition under inflammatory conditions. Next, anti-*Ccne1* siRNA was encapsulated into LNPs and administered to J774A.1 cells to assess the impact of siRNA-mediated Cyclin E1 knockdown. LNPs containing control siRNA served as treatment controls. Cells were pre-incubated with siRNA-loaded LNPs and subsequently polarized as described above (Figure 7A). After harvesting, mRNA expression of major cyclins was analyzed by qPCR (Figure 7D). In M0-polarized J774A.1 cells, treatment with anti-*Ccne1* siRNA LNPs resulted in a significant reduction of *Ccne1* mRNA expression compared to control siRNA-treated cells, confirming efficient knockdown. In addition, expression levels of *Ccna2*, *Ccnb1*, and *Ccnd1* were significantly decreased following anti-*Ccne1* siRNA treatment. M1 polarization itself reduced the expression of these cell cycle-associated genes independently of LNP treatment. Notably, *Ccne2* expression was increased in M1-polarized cells compared to M0 controls but was significantly reduced upon anti-*Ccne1* siRNA LNP treatment relative to control siRNA LNPs.

**Figure 7:**
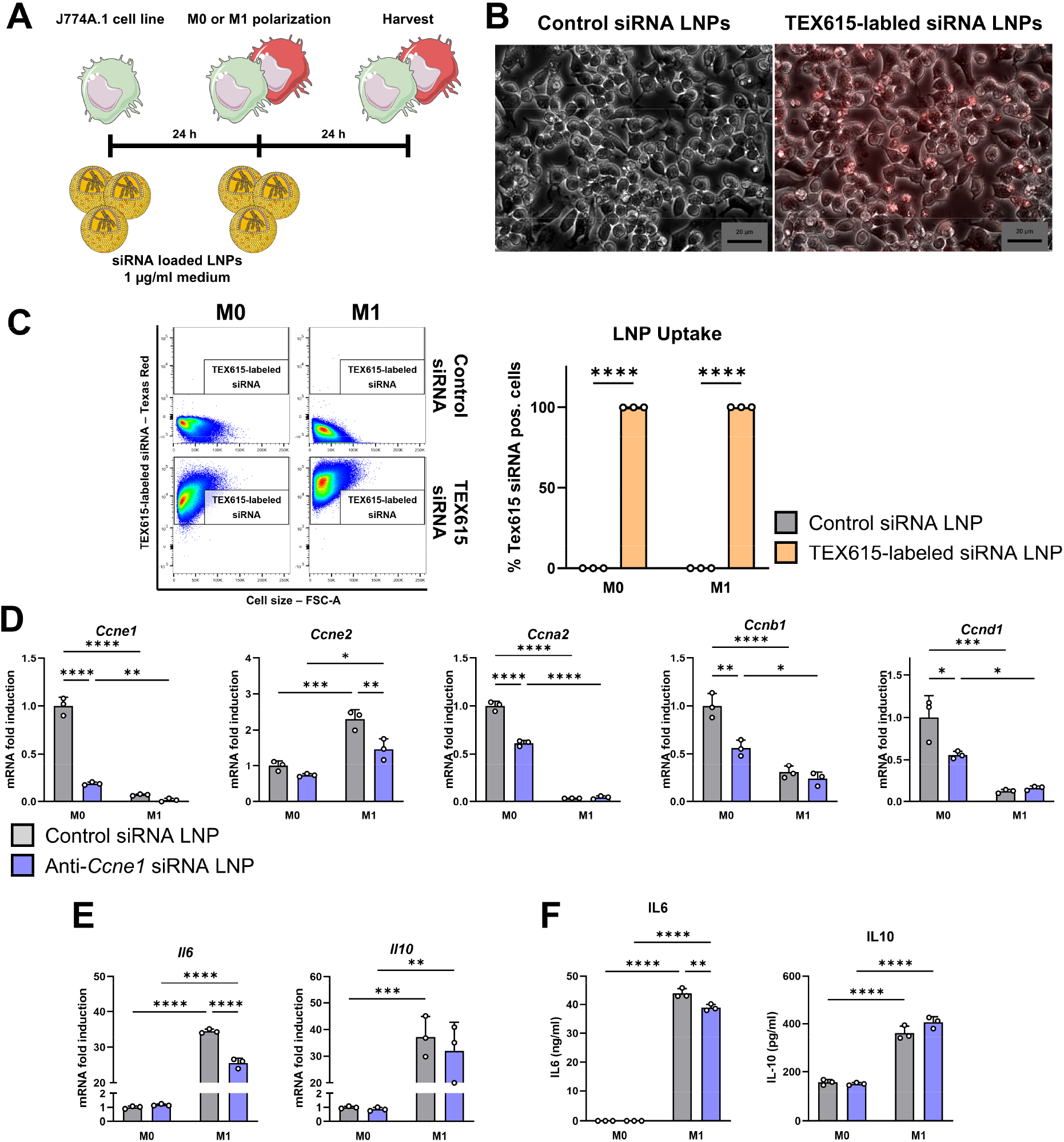
SiRNA-mediated knockdown of *Ccne1* expression reduces *Il6* expression in J774A.1 macrophages The J774A.1 macrophage cell line was utilized to study the impact of lipid nanoparticle (LNP) delivered anti-*Ccne1* siRNA. (**A**) J774A.1 cells were pre-treated with siRNA-loaded LNPs for 24 h followed by M0 or M1 polarization with renewal of siRNA LNPs for another 24 h. (**B**) Control siRNA or TEX615 fluorescently labeled siRNA was encapsulated in LNPs and given to M0 J774A.1 cells. Microscopic images from phase contrast and red fluorescence channels were acquired and are represented as channel overlay images. (**C**) M0 or M1-polarized J774A.1 cells treated with control or TEX615-labeled siRNA delivered by LNPs were analyzed for their uptake efficiency using flow cytometry. Representative dot plots are shown on the left, the quantification from n = 3 per group are shown on the right. M0 or M1-polarized J774A.1 cells were treated with control or anti-*Ccne1* siRNA encapsulated in LNPs (each n = 3). Gene expression for relevant cyclins (**D**) and inflammation-associated mediators (**E**) was determined by qPCR. Gene expression was normalized to *Actb* expression and calculated as fold induction compared to M0 cells treated with control siRNA LNP. (**F**) Protein levels of IL6 and IL10 were determined in cell culture supernatants by ELISA. Data is shown as mean ± SD. *: *p* ≤ 0.05, **: *p* ≤ 0.01, ***: *p* ≤ 0.001, ****: *p* ≤ 0.0001.

To evaluate the functional consequences of siRNA-mediated Cyclin E1 knockdown on inflammatory macrophage responses, *Il6*/IL6 and *Il10*/IL10 expression were analyzed by qPCR and by ELISA in cell culture supernatants (Figure 7E+F). Both cytokines were induced upon M1 polarization at the transcript and protein level, irrespective of the administered siRNA-loaded LNPs. Importantly, M1-polarized J774A.1 cells treated with anti-*Ccne1* siRNA-loaded LNPs displayed reduced *Il6* mRNA expression and decreased IL6 protein secretion compared to cells receiving control siRNA-loaded LNPs. In contrast, *Il10*/IL10 expression was not altered by *Ccne1* knockdown under these conditions.

Collectively, these data demonstrate that J774A.1 macrophages efficiently take up siRNA-loaded LNPs under both M0 and M1 conditions, enabling effective LNP-mediated knockdown of *Ccne1*. Functionally, *Ccne1* knockdown attenuated *Il6*/IL6 expression in M1-polarized J774A.1 cells, supporting the concept that pharmacological targeting of Cyclin E1 may dampen selected pro-inflammatory macrophage responses.

## Discussion

Cyclin E1 is classically known as a regulator of G1/S-phase transition through activation of CDK2 and downstream cell cycle-associated transcriptional programs^10^. However, accumulating evidence indicates that E-type cyclins exert context-dependent functions that extend beyond canonical cell cycle control. While Cyclin E1 and Cyclin E2 are required during embryogenesis, postnatal deletion studies and tissue-specific models have demonstrated that E-type cyclins are not universally essential for S-phase entry in all mammalian cells, likely due to functional redundancy within the cyclin/CDK network^11, 20, 21^. In the liver, Cyclin E1 has emerged as a disease-relevant mediator of regeneration, fibrosis, and hepatocarcinogenesis, with several functions being only partially dependent on CDK2 activity^13–17^. However, most previous studies relied on ubiquitous or systemic Cyclin E1 inhibition and therefore did not resolve whether immune cells contribute to the disease-promoting functions of Cyclin E1. Here, we identify Cyclin E1 as a regulator of hepatic immune cell composition and macrophage-associated inflammatory signaling during acute immune-mediated liver injury.

Together, our findings support a model in which Cyclin E1 promotes acute immune-mediated liver injury through two complementary mechanisms: maintenance of selected hepatic immune cell populations, particularly Kupffer cells, and regulation of inflammatory transcriptional programs in activated macrophages.

### Cyclin E1 shapes the hepatic immune cell compartment

A major finding of this study is that constitutive Cyclin E1 deficiency altered the hepatic immune cell landscape already under basal conditions. *Ccne1*^−/−^ mice displayed reduced numbers of hepatic CD45^+^ immune cells, which was particularly evident for Kupffer cells as well as CD4^+^ and CD8^+^ T cells. This observation was unexpected, as *Ccne1*^−/−^ mice have previously been described as viable and largely without overt developmental abnormalities^11^. Our data therefore suggests that Cyclin E1 may contribute to the establishment or maintenance of selected hepatic immune cell populations, even in the absence of inflammatory challenge.

The reduced number of hepatic T cells may reflect altered recruitment, survival, or expansion within the liver microenvironment. Although the role of E-type cyclins in mature immune cells remains poorly defined, Cyclin E/CDK2-dependent mechanisms have been linked to regulation of FOXP3 and T-cell function^22^. Whether Cyclin E1 directly affects T-cell proliferation or differentiation was not addressed in the present study and remains an important question for future work.

The reduction in Kupffer cell numbers is particularly relevant to the liver phenotype observed in this study. Kupffer cells are self-renewing, liver-resident macrophages derived from embryonic progenitors and are central regulators of hepatic immune tolerance and inflammation^1^. Thus, reduced Kupffer cell abundance in *Ccne1*^−/−^ mice may result from altered embryonic development, impaired niche adaptation, reduced self-renewal, or altered survival. Because bone marrow-derived macrophage differentiation was not impaired by Cyclin E1 deficiency *in vitro*, the Kupffer cell phenotype may reflect a liver-specific requirement for Cyclin E1 in resident macrophage development or maintenance rather than a general defect in myeloid differentiation.

### Macrophage differentiation and cell cycle progression are largely independent of Cyclin E1

Despite altered hepatic immune cell composition *in vivo*, *Ccne1*^−/−^ bone marrow precursor cells differentiated efficiently into BMDMs *in vitro*. Differentiation kinetics, cell cycle phase distribution, and expression of major cell cycle regulators were largely comparable between wildtype and *Ccne1*^−/−^ cells. These findings are consistent with previous observations that Cyclin E1 is dispensable for proliferation in several cell types, including mouse embryonic fibroblasts and myeloid progenitors^11, 12, 23^. They further support the concept that myeloid cell proliferation can proceed independently of Cyclin E1, potentially through compensation by other cyclins or alternative CDK complexes.

Notably, we did not observe compensatory upregulation of *Ccne2*, *Ccna2*, *Ccnb1*, *Ccnd1*, or *Cdk2* during macrophage differentiation. This suggests that the preserved cell cycle activity in *Ccne1*^−/−^ BMDMs is not explained by transcriptional induction of the analyzed cyclins. Instead, compensation may occur through post-transcriptional regulation, altered kinase complex formation, or redundancy within the broader cell cycle machinery. Previous work has shown that Cyclin A-type proteins are particularly important for hematopoiesis, whereas Cyclin E1 expression is comparatively low in bone marrow cells^24^. Thus, Cyclin E1 appears dispensable for macrophage differentiation under the experimental conditions used in our present study.

Bulk RNA-seq nevertheless revealed reduced expression of cell cycle-related gene programs in unstimulated *Ccne1*^−/−^ BMDMs, including decreased inferred activity of E2F transcription factors. This finding indicates that *Ccne1* deficiency may subtly affect basal transcriptional cell cycle networks without producing measurable changes in cell cycle phase distribution by flow cytometry. Such a discrepancy may reflect compensatory mechanisms that maintain cell cycle progression despite reduced expression of cell cycle-related transcriptional programs.

### Cyclin E1 modulates inflammatory macrophage activation

Although Cyclin E1 was dispensable for macrophage differentiation and cell cycle progression, it influenced the inflammatory response of M1-polarized macrophages. Wildtype and *Ccne1*^−/−^ BMDMs both responded efficiently to LPS and IFNγ stimulation, as shown by induction of M1-associated markers. However, *Ccne1*-deficient macrophages displayed reduced *Il6*/IL6 expression and secretion, while *Il10* gene expression showed a trend toward increased levels. These findings argue for a non-canonical role of Cyclin E1 in controlling macrophage inflammatory gene expression rather than a simple effect on macrophage proliferation or polarization efficiency.

IL6 is a central inflammatory cytokine with context-dependent functions in the liver. It contributes to acute phase responses and liver regeneration, but chronic or excessive IL6 signaling is associated with liver inflammation, fibrosis, cirrhosis, and hepatocarcinogenesis^25^. In experimental HCC, IL6 deletion in bone marrow-derived immune cells has been shown to suppress tumor initiation^26^. Previous work from our group further demonstrated that interventional Cyclin E1 deletion in inflammatory HCC reduced hepatic IL6 levels and tumor burden, whereas CDK2 deletion did not reproduce this effect^16^. Together with the present data, these findings support the concept that Cyclin E1 promotes inflammatory signaling through mechanisms that are at least partly independent of canonical Cyclin E1-driven cell cycle progression.

The underlying molecular mechanism remains unresolved. One possibility is that Cyclin E1 directly or indirectly modulates transcriptional activity at inflammatory gene loci such as *Il6*. Non-canonical transcriptional functions have been described for other cyclin family members, including D-type cyclins, which can regulate gene expression independently of their classical kinase partners^27–29^. The RNA-seq data support this concept by showing reduced inflammatory pathway enrichment and decreased inferred activity of IRF1 and NF-κB1 in M1-polarized *Ccne1*^−/−^ BMDMs. Thus, Cyclin E1 may influence inflammatory macrophage activation by modulating NF-κB- or interferon-associated transcriptional networks.

### Cyclin E1 promotes ConA-induced immune-mediated liver injury

The *in vivo* relevance of Cyclin E1-dependent inflammatory regulation was demonstrated in the ConA model of acute immune-mediated hepatitis. *Ccne1*^−/−^ mice were strongly protected from ConA-induced mortality and displayed markedly reduced liver necrosis and lower activation of apoptosis-associated caspase pathways. These findings identify Cyclin E1 as a driver of acute immune-mediated liver injury.

The protective phenotype was associated with altered hepatic immune cell composition and reduced inflammatory mediator expression. In ConA-treated *Ccne1*^−/−^ mice, Kupffer cell numbers remained significantly lower than in wildtype controls, while monocyte and monocyte-derived macrophage numbers were largely unchanged. In parallel, hepatic expression of macrophage-associated cytokines and chemokines, including IL6, TNF, CCL2, and IL10, was reduced. These findings suggest that Cyclin E1 deficiency attenuates ConA-induced inflammation through at least two non-mutually exclusive mechanisms: reduced abundance of liver-resident macrophages and impaired inflammatory cytokine production by macrophages as supported by the BMDM experiments. Similar results were observed in a model of genetically induced chronic hepatitis where overall reduced immune cell numbers and reduced expression of pro-inflammatory TNF were observed in *Ccne1*-deficient mice^30^.

The interpretation of reduced IL6 in the ConA model requires consideration of its stage-dependent functions. Previous studies suggested that IL6 may exert hepatoprotective effects during early phases of ConA-induced injury, while contributing to detrimental inflammatory responses at later stages^31^. Because our analyses were performed 24 hours after ConA administration, the reduced IL6 levels in *Ccne1*^−/−^ mice likely reflect attenuation of a late inflammatory phase. This is consistent with the observed reduction in necrosis, caspase activation, and mortality.

While the ConA model is strongly T cell-driven, macrophages and Kupffer cells amplify inflammatory injury through cytokine and chemokine production. Therefore, the reduced CD8^+^ T cell numbers observed in *Ccne1*^−/−^ livers may also contribute to protection. At present, the relative contribution of macrophages, T cells, hepatocytes, and other non-parenchymal cells cannot be fully separated because a constitutive knockout model was used. Moreover, systemic effects of *Ccne1* deficiency, including potential alterations in the gut-liver axis, cannot be excluded. Zhou *et al.* recently reported that ConA disrupts intestinal barrier integrity and promotes bacterial translocation to the liver^32^, which could be modulated due to loss of Cyclin E1 in intestinal cells.

### siRNA-mediated Cyclin E1 knockdown attenuates IL6 expression in macrophage-like cells

A translationally relevant aspect of this study is the demonstration that siRNA-loaded lipid nanoparticles can efficiently deliver anti-*Ccne1* siRNA into J774A.1 macrophage-like cells. Cyclin E1 knockdown reduced *Il6*/IL6 expression in M1-polarized J774A.1 cells, thereby reproducing a central feature of the inflammatory phenotype observed in constitutive *Ccne1*^−/−^ BMDMs. Together with previous findings showing that siRNA-mediated Cyclin E1 inhibition attenuates experimental liver fibrosis and inflammation *in vivo*^17^, these results support the potential of Cyclin E1 as a pharmacologically accessible regulator of inflammatory liver disease.

The effects of anti-*Ccne1* siRNA were not restricted to *Ccne1* but also included reduced expression of several other cyclins. Because the siRNA sequence was selected for specificity toward *Ccne1*, these changes may reflect secondary alterations in cell cycle-associated transcriptional programs rather than direct off-target effects. Importantly, the reduction in *Il6*/IL6 was observed under pro-inflammatory conditions, supporting a functional connection between *Ccne1* expression and macrophage inflammatory activity. However, because J774A.1 cells are immortalized and proliferative, the magnitude of this response may differ from that in primary macrophages.

Overall, the LNP experiments provide proof of principle that acute pharmacological reduction of Cyclin E1 can attenuate selected pro-inflammatory macrophage responses. Although efficient LNP uptake in cultured macrophages does not establish macrophage-selective delivery *in vivo*, the present findings complement the protective phenotype of *Ccne1*^−/−^ mice and support further evaluation of Cyclin E1 inhibition as an immunomodulatory approach in inflammatory liver disease.

### Limitations and future directions

This study has several limitations. First, the use of constitutive *Ccne1*^−/−^ mice precludes definitive assignment of the protective ConA phenotype to a specific cell type. *Ccne1* deficiency affects not only macrophages but also hepatocytes, hepatic stellate cells, endothelial cells, and lymphoid populations. Thus, macrophage-specific or Kupffer cell-specific *Ccne1* deletion models will be required to determine whether Cyclin E1 acts directly in hepatic macrophages during acute hepatitis.

Second, analyses performed at the 24-hour endpoint were restricted to surviving animals and may therefore be affected by survivor bias. The surviving wildtype mice may represent a less severely affected subgroup, potentially leading to an underestimation of genotype-dependent differences in liver injury and inflammation. Although the cause of death could not be determined directly, the marked liver injury observed after ConA administration suggests that mortality was associated with severe immune-mediated acute liver failure.

Third, the molecular mechanism linking Cyclin E1 to *Il6* expression remains unresolved. The RNA-seq data suggest reduced inflammatory transcription factor activity, particularly involving IRF1 and NF-κB-associated pathways, but direct regulation has not been demonstrated. Future chromatin-based and proteomic approaches may help clarify whether Cyclin E1 interacts with inflammatory transcriptional complexes.

Finally, the therapeutic potential of Cyclin E1 inhibition in macrophage-driven liver inflammation remains to be tested *in vivo* using targeted LNP delivery. Such studies should evaluate efficacy, biodistribution, immune cell specificity, and safety, particularly given the context-dependent roles of Cyclin E1 and IL6 in liver regeneration, inflammation, and tumorigenesis.

## Conclusion

In summary, this study identifies Cyclin E1 as a previously unrecognized regulator of hepatic immune responses and macrophage-associated inflammatory signaling. Cyclin E1 deficiency altered the hepatic immune cell compartment, reduced Kupffer cell abundance, attenuated macrophage inflammatory cytokine production, and protected mice from ConA-induced acute liver injury. Importantly, Cyclin E1 was dispensable for BMDM differentiation and major cell cycle progression, suggesting that its pro-inflammatory function in macrophages is at least partly independent of canonical cell cycle regulation. Finally, siRNA-mediated *Ccne1* knockdown reduced IL6 expression in macrophage-like cells, supporting the concept that Cyclin E1 may represent a therapeutically targetable regulator of inflammatory liver disease.

## Materials and Methods

### Animal experiments

C57BL/6 mice with constitutive knockout of Cyclin E1 (referred to as *Ccne1*^−/−^) were kindly provided by Piotr Sicinski (Dana-Farber Cancer Institute, Boston, USA) and bred from both, homozygous and heterozygous breeding couples. Mice were housed under controlled environmental conditions (12 h light/dark cycles, constant temperature and humidified atmosphere) with *ad libitum* water and food at the Institute of Laboratory Animal Science of the Uniklinik RWTH Aachen.

Wildtype littermates were used as controls. For characterization of homeostatic hepatic immune cell composition in wildtype and *Ccne1*^−/−^ mice and for the Concanavalin A mouse model, male 8 - 10-week-old mice were used. Mice received an initially considered sublethal dose of 20 mg/kg ConA (Sigma Aldrich, St. Louis, USA) dissolved in 0.9 % NaCl by tail vein injection and were sacrificed for analysis after 24 h.

All animal experiments were approved by the Ministry for Environment, Agriculture, Conservation and Consumer Protection of the State of North Rhine-Westphalia (LANUV) and were carried out according to the German legal requirements.

### Isolation, culture and polarization of bone marrow-derived cells

Bone marrow cell isolation and culture were performed as previously described with minor modifications^33^. In brief, bone marrow was isolated from femur and tibia of 8–12-week-old wildtype or *Ccne1*^−/−^ mice and cultured in RPMI 1640 containing 10 % fetal calf serum, 1 % penicillin/streptomycin and 20 % L929 conditioned medium for three days (37 °C, 5 % CO_2_). Cells were harvested and seeded into six-well plates at a density of 5 x 10^5^ cells per well and cultured for additional three days to obtain fully differentiated bone marrow derived macrophages.

Pro-inflammatory polarization of BMDMs (referred to M1) was induced by cell culture medium supplementation with 100 ng/ml LPS and 100 ng/ml IFNγ for 24 h; anti-inflammatory polarization (referred to M2a) was induced by supplementation with 20 ng/ml IL4 and 20 ng/ml IL13 for 24 h. Non-polarized cells (referred to as M0) were used as control.

To investigate bone marrow cells during differentiation, 1.5 x 10^6^ cells of freshly isolated bone marrow cells were seeded per well, and samples were collected and analyzed on six consecutive days.

### Culture and siRNA-LNP treatment of J774A.1 cell line

The J774A.1 cell line was bought from Sigma Aldrich (St. Louis, USA) and cultured in DMEM containing 10 % FCS and 1 % penicillin/streptomycin at 37 °C and 5 % CO_2_. Cells were regularly tested for mycoplasma contamination.

SiRNAs (Supplementary Table 1) were obtained from Integrated DNA Technologies (Coralville, USA) and encapsulated in LNPs containing Dlin-MC3-DMA (Hycultec, Beutelsbach, Germany) as recently described^34^. Particle size was determined by dynamic light scattering and was on average 86 ± 14 nm.

5 x 10^5^ J774A.1 cells were seeded per well in six-well plates and cultured for three days prior to addition of 2 µg encapsulated siRNA per well. After 24 h medium was renewed with addition of fresh siRNA LNPs as well as LPS and IFNγ to induce M1 polarization as described above. Cells were harvested and subsequently analyzed after 24 h.

### Histopathology

Liver tissues were dissected and fixed in 4 % paraformaldehyde for 24 h. Dehydration, paraffin embedding, slicing and hematoxylin and eosin staining (H&E) was performed by the Immunohistochemistry Facility of the Interdisciplinary Center for Clinical Research (IZKF) of the RWTH Aachen by standard methods. Bright field images of whole liver sections were captured on a Zeiss Axio Observer.Z1 microscope equipped with a motorized stage followed by shading correction and tile stitching using Zen Pro software v2.6. Necrotic tissue area fractions were quantified using Fiji v1.52s by applying color thresholding to determine necrotic area and comparing it to the whole section area.

### Flow cytometry

Liver lobes were minced and digested with 1.6 mg/ml collagenase IV (Worthington, Lakewood, USA) and 50 µg/ml DNase I (Roche, Basel, Switzerland) dissolved in DMEM by shaking for 45 min at 37 °C and 150 rpm. Digested tissue was filtered and washed through a 70 µm cell strainer with HBSS containing 1.5 mM EDTA (AppliChem, Darmstadt, Germany) and 0.06 % bovine serum albumin followed by centrifugation at 50 x g for 5 min to separate hepatocytes. Supernatants were collected by centrifugation at 300 x g for 10 min and red blood cells were lysed with RBC lysis buffer (eBioscience, San Diego, USA). Cells were collected in FACS tubes and stained, measured and analyzed as recently described^16^. Applied gating strategies are shown in Supplementary Figure 1, and details of used antibodies can be found in Supplementary Table 2.

### Western blot analysis

Protein isolation and Western blot analysis was conducted as previously described^35^. Used antibodies are listed in Supplementary Table 3. Detection of GAPDH was used as loading control. Blots were visualized by ECL Prime Western-Blot-System (Cytiva, Marlborough, USA) using an ImageQuant LAS 4000 System with LAS 4000 v1.2 Control Software (GE Healthcare, Chicago, USA) according to the manufacturer’s protocol. Uncropped Western blots can be found in Supplementary Figure 5.

### Enzyme-linked immunosorbent assay (ELISA)

Concentrations of IL6, TNF, IFNG, IL10 and CCL2 in cell culture supernatant and in liver protein extracts were determined using Ready-SET-Go! Mouse ELISA Kits (eBioscience, San Diego, USA) following manufacturer’s instructions and a Cytation 3 micro well reader with Gen5 v3.00.19 software (Biotek Instruments, Winooski, USA). For liver tissue analyses, cytokine and chemokine concentrations were normalized to the total protein concentration determined by Bradford assay.

### RNA isolation

Total RNA was isolated from cells and liver tissue using QIAzol lysis reagent (Qiagen, Hilden, Germany) following manufacturer’s instructions with modified precipitation in isopropanol overnight at −20 °C.

### Quantitative real-time PCR (qPCR)

2 µg RNA were transcribed into cDNA using the Omniscript RT Kit and oligo-dT primers following manufacturer’s instructions (Qiagen, Hilden, Germany). QPCR was performed using Fast SYBR Green Mastermix (Applied Biosystems, Waltham, USA) on a QuantStudio 6 Flex PCR system. Used primer sequences are listed in Supplementary Table 4. Analysis was performed using the 2^−Δ^ ^ΔCt^-method^36^ with *Actb* as housekeeping gene for expression normalization. Fold changes were calculated as indicated in the respective figure legends.

### Bulk RNA sequencing

RNA libraries were prepared using the Collibri 3’mRNA Library Preparation Kit (Invitrogen, Carlsbad, USA). Libraries were sequenced on a NextSeq 500 System (Illumina) followed by read alignment to the GRCm39 reference genome using STAR v2.7.11b^37^ and expression quantification with Salmon v1.10.3^38^ using the nf-core/rnaseq pipeline v3.26.0^39^. Normalization and differential gene expression analysis was performed in R v4.5.1 using DESeq2 v1.48.2^40^. Genes with *p* < 0.05 after Benjamini-Hochberg correction were considered as significantly differentially expressed. Overrepresentation analysis on gene ontology pathways was performed using the clusterProfiler package v4.16.0^41^ (cutoff: adj. *p* < 0.1). We used the decoupleR package v2.14.0^42^ together with the CollecTRI network to infer transcription factor activity scores from our expression data.

### Statistical analysis

Statistical analyses were performed using GraphPad Prism 11. Data are presented as mean ± SD unless indicated otherwise. Survival proportions after 24 h were analyzed using Fisher’s exact test. For comparisons between two groups, normality was assessed using the Shapiro-Wilk test. Non-normally distributed data were analyzed using the Mann-Whitney U test. Normally distributed data were analyzed using an unpaired two-tailed Student’s t test or, in case of unequal variances as determined by the F test, an unpaired t test with Welch’s correction.

For comparisons of more than two groups with one experimental variable, one-way ANOVA followed by Tukey’s multiple comparisons test was used.

For experiments with two independent variables, ordinary two-way ANOVA was applied, followed by Tukey’s multiple comparisons test. Outliers were identified using Grubbs’ test with α = 0.05 and excluded where applicable. Statistical significance was defined as *p* ≤ 0.05.

## Data availability

The RNA-sequencing data and other data supporting the findings of this study are available from the corresponding author upon reasonable request.

## Declaration of generative AI in the manuscript preparation process

During the preparation of this work, the authors used Microsoft Copilot to improve language and readability. After using this tool, the authors reviewed and edited the content as needed and took full responsibility for the content of the published article.

## Acknowledgements

This work was supported by the Immunohistochemistry and Genomics Facility of the Interdisciplinary Center for Clinical Research (IZKF) within the Faculty of Medicine at RWTH Aachen University. Illustrations in Figure 2A, 5A, 7A and Supplementary Figure 2A were created using images provided by Servier Medical Art (https://smart.servier.com) licensed under CC BY 4.0 (https://creativecommons.org/licenses/by/4.0/). This work was supported by the German Research Foundation (Deutsche Forschungsgemeinschaft, DFG) through CRC 1382 “Gut-Liver Axis” (Projects A02, B05, project number 403224013 to C.L. and F.T.), RTG 2375 “Tumor-Targeted Drug Delivery” (project number 331065168 to C.P. and C.L.), CRC/TR 412 (project number 535081457 to F.T.), and the Germany’s Excellence Strategy through EXC 3118/1 (project number 533770413 to F.T.). M.B. received funding from the Leibniz ScienceCampus ACTISONO, supported by the Leibniz Association (grant number W89/2023), and from the DFG (project number 417882771).

## Author contributions

Christian Penners: Conceptualization, Data Curation, Formal Analysis, Investigation, Methodology, Software, Validation, Visualization, Writing – Original Draft, Writing – Review & Editing.

Cheng Lin: Investigation, Methodology, Writing – Review & Editing.

Alexander M. Jans: Investigation, Methodology, Writing – Review & Editing.

Antje Mohs: Investigation, Methodology, Writing – Review & Editing.

Frank Tacke: Conceptualization, Funding Acquisition, Supervision, Writing – Review & Editing. Matthias Bartneck: Funding Acquisition, Methodology, Resources, Writing – Review & Editing. Roland Sonntag: Conceptualization, Data Curation, Formal Analysis, Investigation, Project Administration, Resources, Supervision, Validation, Writing – Original Draft, Writing – Review & Editing.

Christian Liedtke: Conceptualization, Formal Analysis, Funding Acquisition, Project Administration, Resources, Supervision, Validation, Writing – Original Draft, Writing – Review & Editing.

## Ethical Statement

The animal study was approved by authority for environment conservation and consumer protection of the state of North Rhine-Westphalia, Recklinghausen, Germany (LANUV). The study was conducted in accordance with the local legislation and institutional requirements.

## Competing interest statement

The authors declare no competing interests.

## Funding

This work was supported by the German Research Foundation (Deutsche Forschungsgemeinschaft, DFG) through CRC 1382 “Gut-Liver Axis” (Projects A02, B05, project number 403224013 to C.L. and F.T.), RTG 2375 “Tumor-Targeted Drug Delivery” (project number 331065168 to C.P. and C.L.), CRC/TR 412 (project number 535081457 to F.T.), and the Germany’s Excellence Strategy through EXC 3118/1 (project number 533770413 to F.T.). M.B. received funding from the Leibniz ScienceCampus ACTISONO, supported by the Leibniz Association (grant number W89/2023), and from the DFG (project number 417882771).

## Supplementary Figures

**Supplementary Figure 1.**
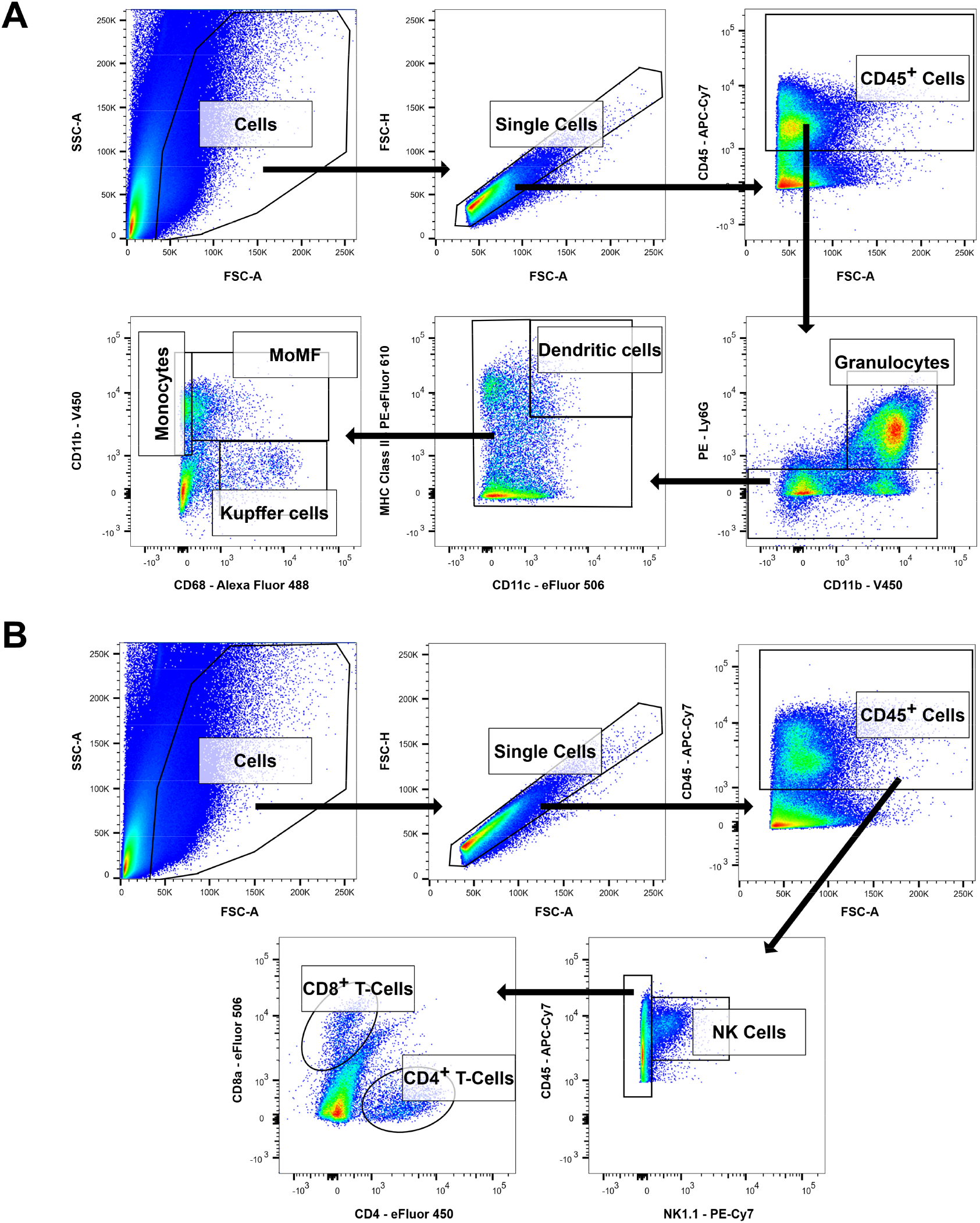
(**A**) Gating strategy to identify lymphoid cell populations: cells with low size (FSC-A) and doublets (FSC-A vs. FSC-H) were excluded from the analysis. Immune cells (CD45^+^) were selected for further analysis. Due to poor staining of the CD3 - PE Antibody (T-cell receptor), this marker was not included in the analysis. Natural killer cells and natural killer T-cells (both referred to as NK-cells, CD45^+^, NK1.1^+^) were gated from total immune cells. The remaining cells were gated for CD4^+^, CD8^−^ cells (referred to as CD4^+^ T-cells) and CD4^−^, CD8^+^ cells (referred to as CD8^+^ T-cells). (**B**) Gating strategy to identify myeloid cell populations: Cells with low size (FSC-A) and doublets (FSC-A vs. FSC-H) were excluded from the analysis. CD45^+^ immune cells were selected for further analysis followed by gating for granulocytes (CD45^+^, CD11b^+^, Ly6G^+^) and dendritic cells (CD45^+^, Ly6G^−^, CD11c^+^, MHC Class II^+^). The remaining cells were defined as monocytes (CD11b^+^, CD68^−^), monocyte-derived macrophages (MoMF, CD11b^high^, CD68^+^) or Kupffer cells (CD11b^low-intermediate^, CD68^+^).

**Supplementary Figure 2.**
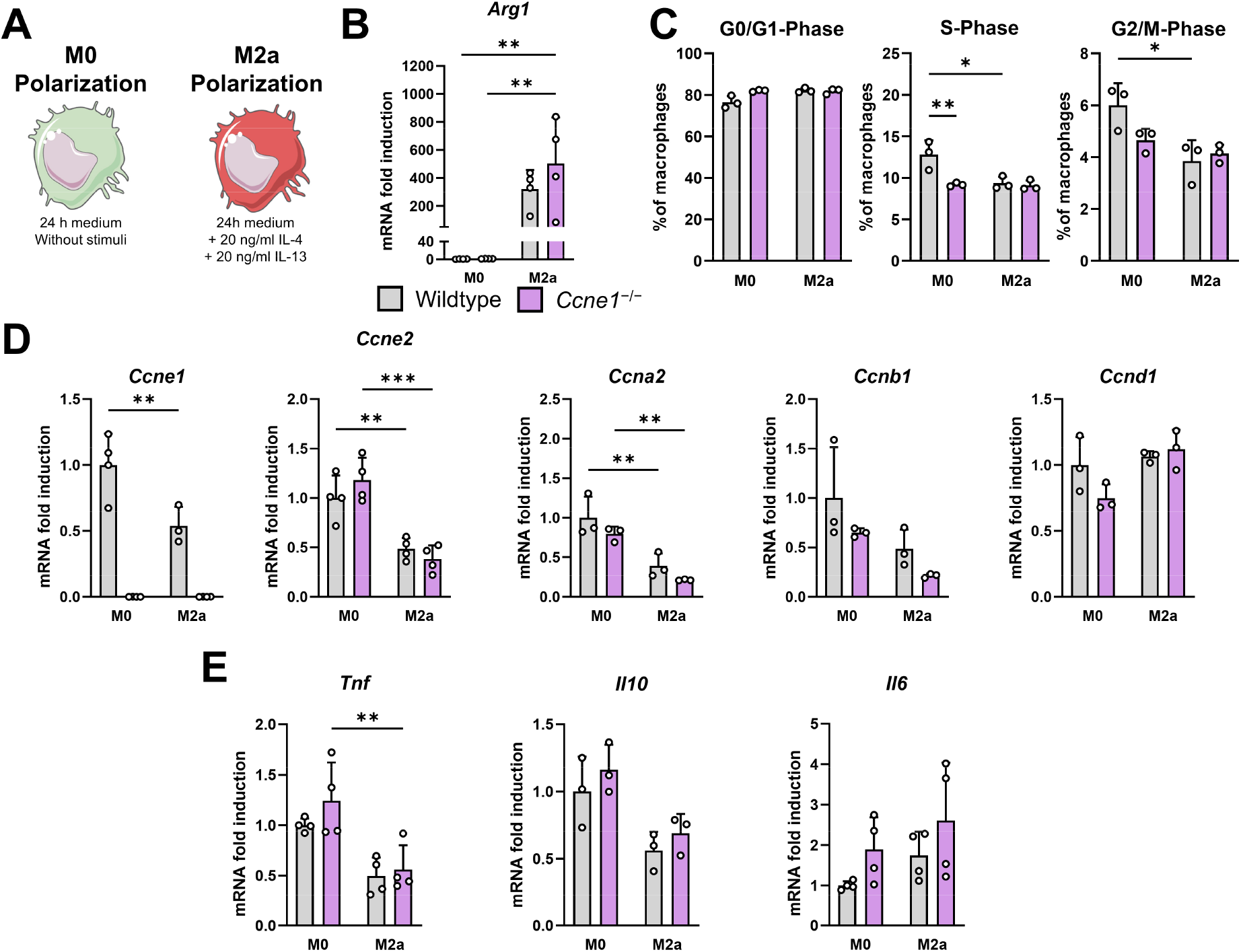
Wildtype (n = 3 - 4) and *Ccne1*^−/−^ (n = 3 - 4) mice were sacrificed to isolate bone marrow for *in vitro* generation of BMDMs followed by their anti-inflammatory polarization (M2a) or were left untreated (M0). (**A**) Schematic representation of M0 or M2a polarization conditions. (**B**) Relative *Arg1* gene expression, a well-established marker for M2a macrophages, was determined by qPCR. Expression was normalized to *Actb* levels and calculated as fold change compared to M0-polarized wildtype BMDMs. (**C**) Cell cycle phases were determined by flow cytometry using DAPI to measure cellular DNA content. The cell numbers in each phase are shown relative to all analyzed BMDMs. Gene expression of cell cycle-related genes (**D**) and inflammatory mediators (**E**) was determined by qPCR. Expression was normalized to *Actb* levels and calculated as fold change compared to M0-polarized wildtype BMDMs. Data is shown as mean ± SD. *: *p* ≤ 0.05, **: *p* ≤ 0.01, ***: *p* ≤ 0.001.

**Supplementary Figure 3.**
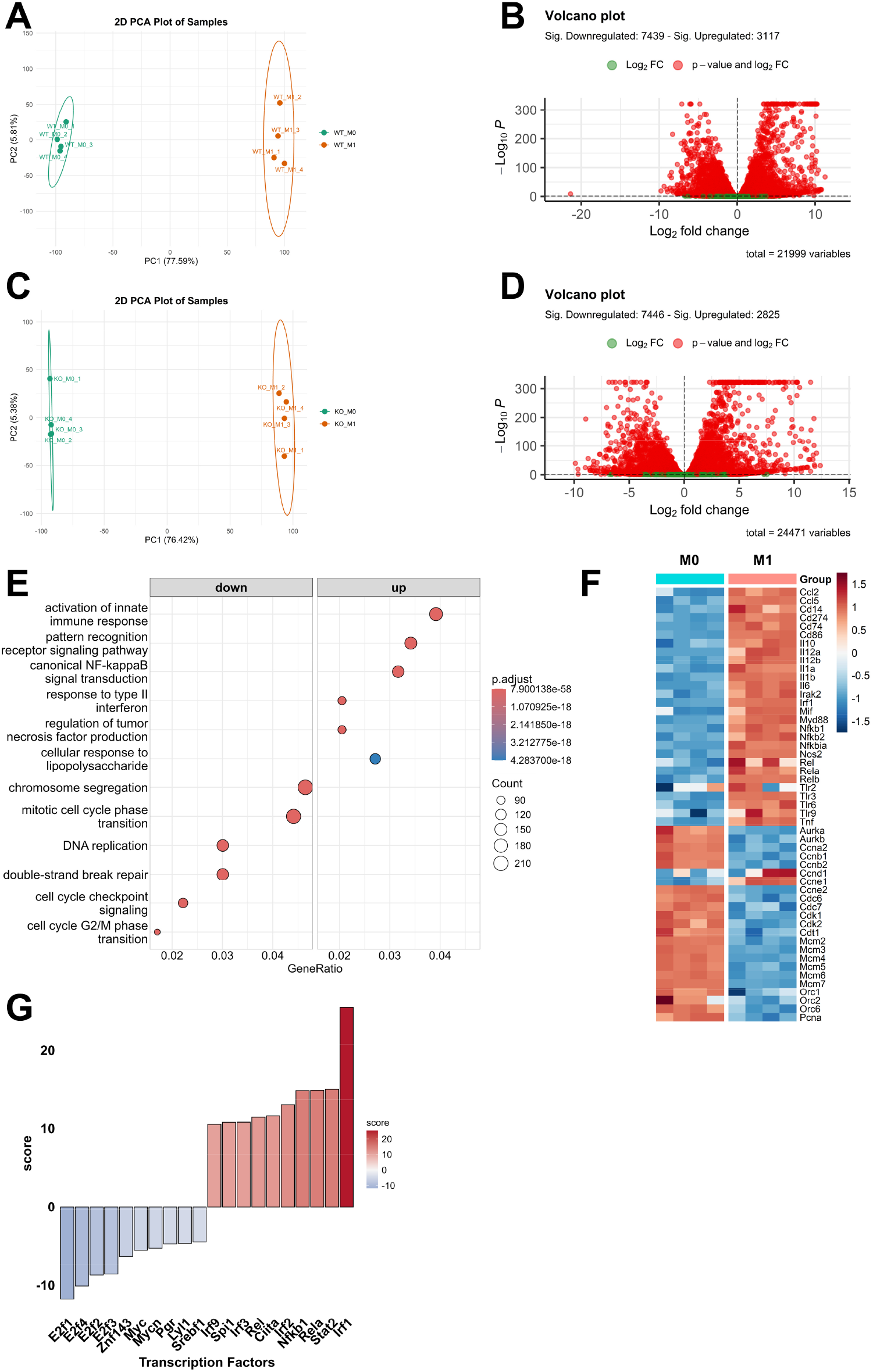
Wildtype (n = 4) and *Ccne1*^−/−^ (n = 4) mice were sacrificed to isolate bone marrow for *in vitro* generation of BMDMs followed by their pro-inflammatory polarization (M1) or were left untreated (M0) as described in Figure 5A. Total mRNA was isolated and bulk RNA-seq was performed. (**A**) Differential gene expression between M1-polarized wildtype BMDMs compared to M0 wildtype BMDMs was analyzed. Principal component analysis (PCA) was performed, and results are shown as PCA plot. (**B**) Volcano Plot showing differential gene expression results with green dots representing non-significant and red dots representing significant deregulated genes (adjusted p-value < 0.05). (**C**) Differential gene expression between M1-polarized *Ccne1*^−/−^ BMDMs compared to M0 *Ccne1*^−/−^ BMDMs was analyzed. Results of principal component analysis (PCA) are shown as PCA plot and results of differential gene expression are shown as volcano plot (**D**). Over representation analysis was performed on up- or down-regulated genes using gene ontology pathways. (**E**) Results of selected significant deregulated pathways are shown as dot plot indicating number of regulated genes by size and statistical significance by color. (**F**) Heatmap of selected genes involved in regulation of inflammation or cell cycle. (**G**) Transcription factor activity scores were calculated using transcription factor activity inference. The ten statistically significant most active or deactivated transcription factors are shown as bar plot.

**Supplementary Figure 4.**
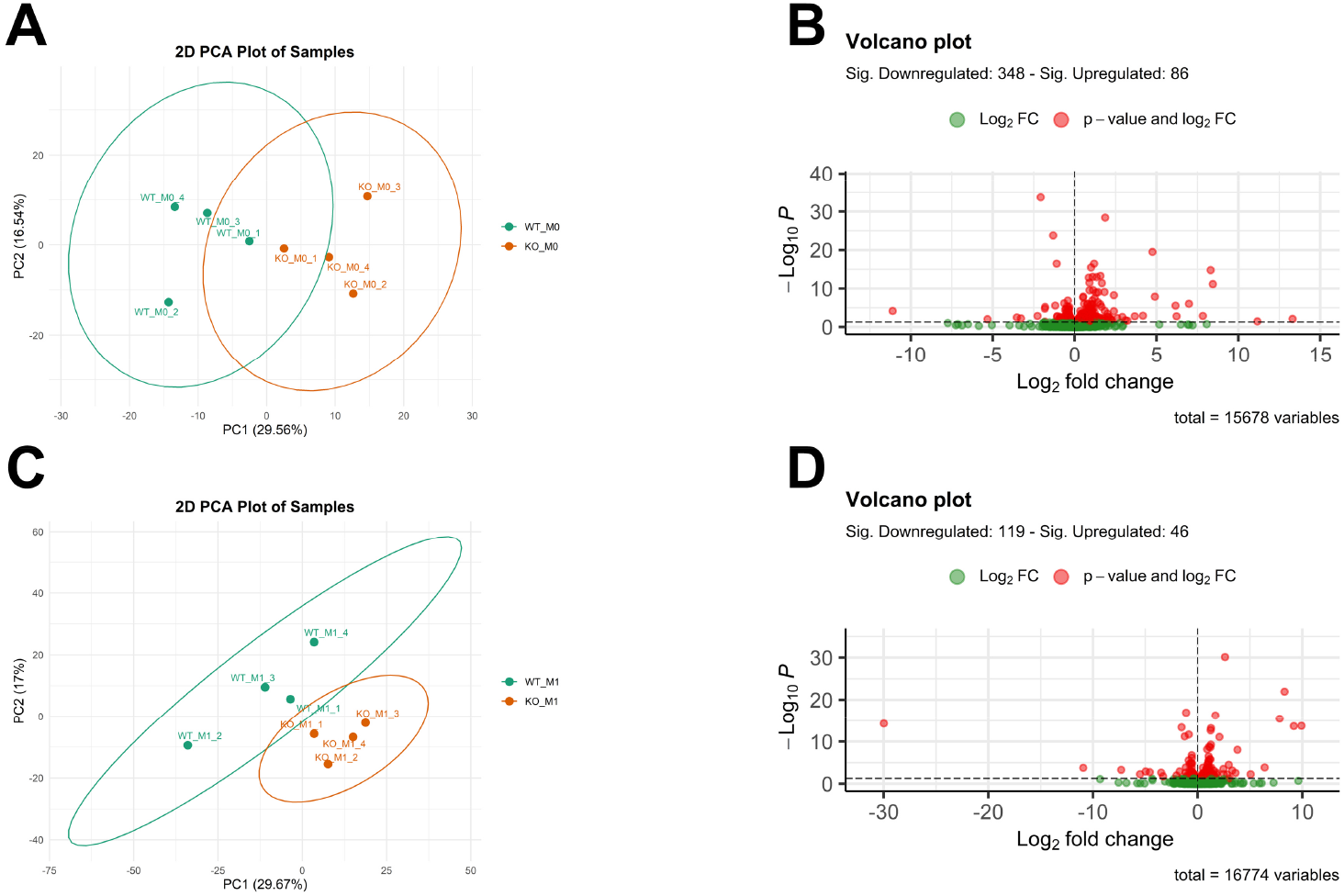
Wildtype (n = 4) and *Ccne1*^−/−^ (n = 4) mice were sacrificed to isolate bone marrow for *in vitro* generation of BMDMs followed by their pro-inflammatory polarization (M1) or were left untreated (M0) as described in Figure 5A. Total mRNA was isolated and bulk RNA-seq was performed. (**A**) Differential gene expression between M0-polarized *Ccne1*^−/−^ BMDMs compared to M0 wildtype BMDMs was analyzed. Principal component analysis (PCA) was performed, and results are shown as PCA plot. (**B**) Volcano Plot showing differential gene expression results with green dots representing non-significant and red dots representing significant deregulated genes (adjusted p-value < 0.05). (**C**) Differential gene expression between M1-polarized *Ccne1*^−/−^ BMDMs compared to M1 wildtype BMDMs was analyzed. Results of principal component analysis (PCA) are shown as PCA plot and results of differential gene expression are shown as volcano plot (**D**).

**Supplementary Figure 5.**
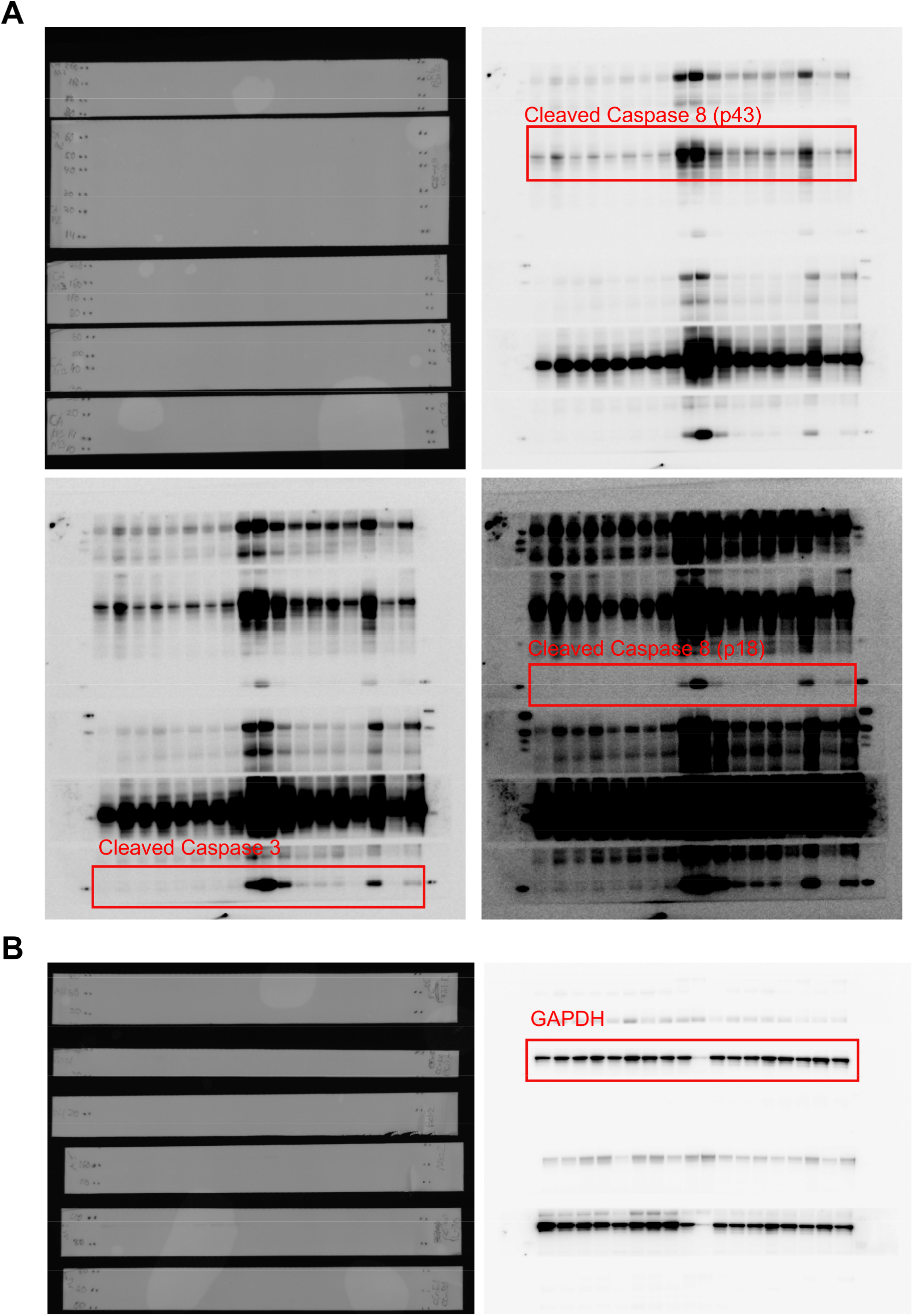
(**A**) Uncropped Western blot membranes for cleaved Caspase-3 and -8 and (**B**) GAPDH.

## Supplementary Tables

**Supplementary Table 1:**
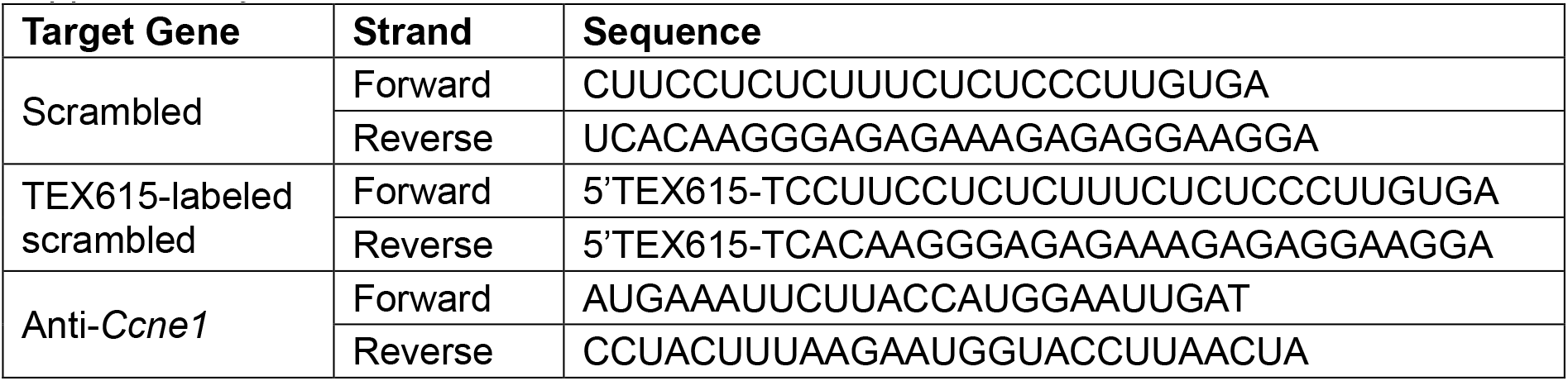
siRNAs.

**Supplementary Table 2:** Antibodies used for flow cytometry.

| Antigen | Fluorophore | Article Number | Company |
| --- | --- | --- | --- |
| CD11b | PE | 12-0112-83 | eBioscience |
| CD11b | V450 | 560455 | BD Biosciences |
| CD11c | eFluor 506 | 69-0114-82 | eBioscience |
| CD4 | eFluor 506 | 48-0041-82 | eBioscience |
| CD45 | APC-Cy7 | 557659 | BD Biosciences |
| CD68 | Alexa Fluor 488 | 53-0681-82 | eBioscience |
| CD68 | eFluor 660 | 50-0681-82 | eBioscience |
| CD8a | eFluor 506 | 69-0081-82 | eBioscience |
| Ly6G | PE | 12-9668-82 | eBioscience |
| MHC Class II | PE-eFluor 610 | 61-5321-82 | eBioscience |
| NK1.1 | PE-Cy7 | 25-5941-81 | eBioscience |

**Supplementary Table 3:**
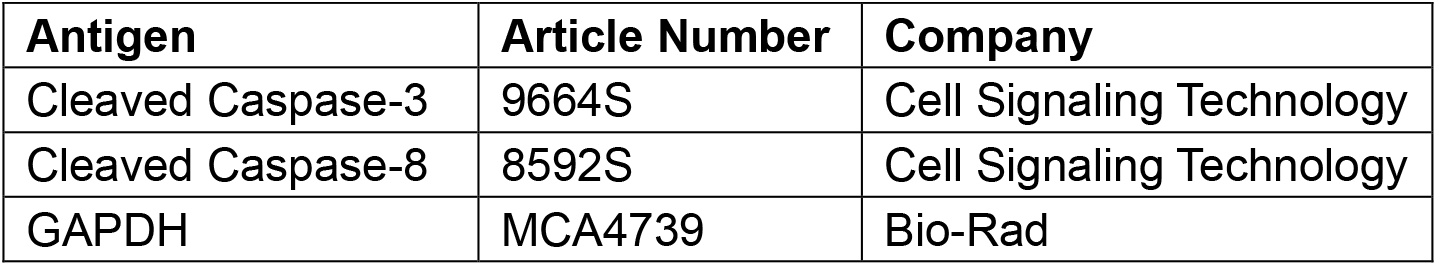
Western blot antibodies.

**Supplementary Table 4:**
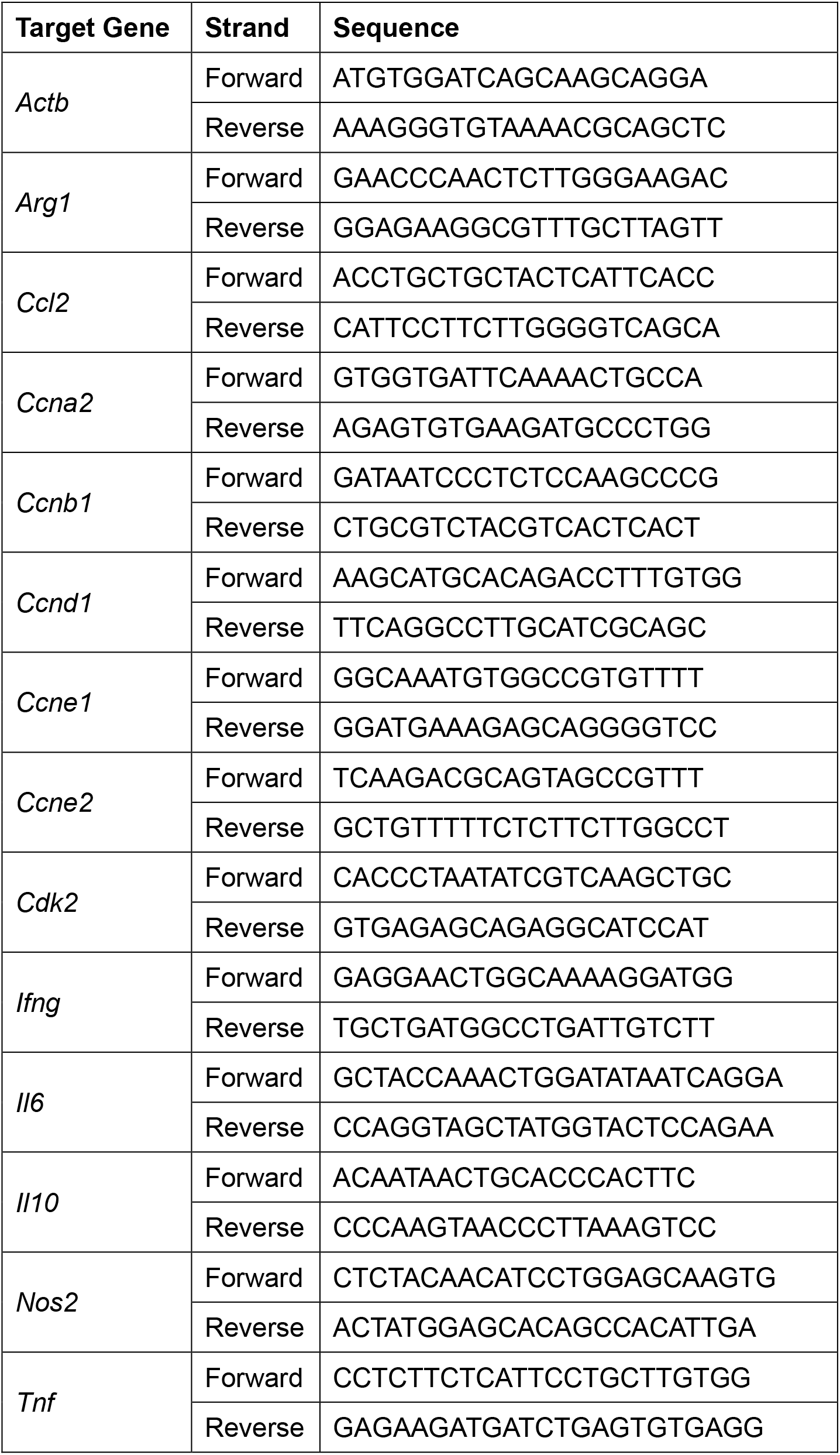
qPCR primer.

## Notes

### Competing Interest Statement

The authors have declared no competing interest.

